# Cellular responsiveness as a predictive indicator for population collapse and autonomous control in continuous cultures of *Pseudomonas putida*

**DOI:** 10.64898/2026.06.08.730862

**Authors:** Maximilian Sehrt, Hannah Sehrt, Laurie Josselin, Juan Andres Martinez, Frédéric Francis, Frank Delvigne

## Abstract

How microbial populations respond to repeated environmental transitions determines both their ecological fitness and their utility in biotechnological applications. Using *Pseudomonas putida* KT2440 equipped with fluorescent biosensors and monitored by automated flow cytometry in the Segregostat platform, we show that exposure to benzoate, a plastic-derived aromatic feedstock, progressively reduces cellular responsiveness, defined as the fraction of cells that successfully activate a gene circuit following an environmental transition. Unlike classical switching costs, which promote phenotypic diversification, benzoate suppresses responsiveness without increasing population entropy, in a concentration-dependent and circuit-independent manner tightly correlated with fitness loss. A resource allocation model incorporating the competing demands of benzoate assimilation, toxicity, and tolerance reveals that this impairment emerges from a three-way competition for limited cellular resources. Above a critical benzoate load, insufficient resources remain available to sustain the adaptive reallocation required for circuit activation. In continuous culture, a non-responsive subpopulation accumulates as a leading indicator of population collapse. Exploiting this signal, we implement a two-stage connected bioreactor system in which benzoate feeding is autonomously regulated based on real-time population structure, enabling complete substrate consumption and stable operation at otherwise destabilizing concentrations. These results establish cellular responsiveness as a quantitative population variable and demonstrate that structure-aware feedback control, acting on population composition rather than bulk physiology, provides a principled route toward autonomous bioprocesses on challenging substrates.

## Introduction

When microbial cells encounter a shift in their chemical environment, they must rapidly reprogram their physiology. This reprogramming relies on the timely activation or repression of specific gene circuits that enable substrate assimilation, stress mitigation, and resource reallocation^1^ ^2^. Although the molecular architecture of many transition-relevant gene circuits has been thoroughly characterized, particularly in the context of diauxic shifts and carbon catabolite repression ^3^ ^4^ ^5^, the factors that determine whether a population engages the required metabolic program rapidly or fails to do so remain poorly understood, especially when the transition involves a metabolically demanding or inhibitory substrate.

Environmental transitions do not elicit uniform responses across a population. Cells within an isogenic culture can adopt distinct response strategies that differ in timing and associated fitness penalties ^6^. We have previously shown that the fitness cost of switching between phenotypic states is a key determinant of population dynamics i.e., high switching costs drive phenotypic diversification, generating coexisting subpopulations rather than a single converged state ^7^. In this context, diversification serves as a population-level buffering strategy, distributing fitness costs across a range of phenotypic states. The Segregostat platform, a cell-machine interface coupling continuous cultivation with automated flow cytometry, has proven central to quantifying and controlling these diversification dynamics in real time ^7^ ^8^ ^9^.

At the single-cell level, the dynamics of gene circuit activation during an environmental transition can be formally described by the concept of first passage time (FPT) i.e., the time elapsed before an individual cell first crosses a defined threshold in gene expression following the transition ^10^ ^11^. FPT distributions capture cell-to-cell variability in activation kinetics and are shaped by both the regulatory architecture of the circuit and the physiological state of the cell upon transition. However, in population-level experiments, where single-cell trajectories are not continuously tracked, FPT cannot be measured directly. We therefore define cellular responsiveness as the population-level analogue of FPT i.e., the fraction of cells in a population that successfully activate a gene circuit above a defined threshold within a fixed time window following an environmental transition. This metric is directly accessible from flow cytometry data, scales naturally with the underlying single-cell FPT distribution and provides a quantitative readout of the population collective capacity to engage a new physiological state.

A distinct and underexplored scenario arises when the environmental transition involves a substrate that not only imposes a fitness cost, but actively impairs the regulatory machinery required for adaptation. Benzoate, a metabolically demanding aromatic compound and an emerging feedstock derived from plastic waste streams ^12^ ^13^ ^14^, presents exactly this challenge in *Pseudomonas putida* KT2440. Benzoate degradation is subject to carbon catabolite repression mediated by the global regulator Crc, which post-transcriptionally represses the *benR* and *benA* transcripts, delaying circuit activation ^15^ ^16^. At elevated concentrations, benzoate additionally imposes metabolic stress that is expected to divert cellular resources toward tolerance and detoxification, leaving fewer resources available for growth and adaptive reprogramming ^17^ ^18^. Whether this dual challenge i.e., regulatory repression combined with resource competition, generates a qualitatively different population response than classical switching costs, and specifically whether it reduces cellular responsiveness rather than promoting diversification, has not been investigated.

Resource allocation models provide a principled framework for interpreting these competing demands. By partitioning cellular biomass among functional sectors i.e., ribosomes, metabolic enzymes, stress-response machinery, these models show how substrate-driven allocation strategies give rise to emergent growth and expression phenotypes ^1^ ^2^ ^19^. Interestingly, Balakrishnan *et al.* demonstrated experimentally in *E. coli* that suboptimal resource allocation during carbon source transitions directly constrains both growth recovery and gene circuit activation, providing a mechanistic link between resource competition and impaired responsiveness. More recent models have extended this framework to antibiotic tolerance ^20^ and to *P. putida* physiology under nutrient limitation ^21^ ^22^. However, no model has yet addressed a substrate that simultaneously demands investment in assimilation machinery, imposes toxicity, and requires a dedicated tolerance sector, conditions that create three-way competition for a limited resource pool and that we hypothesize underlie the loss of responsiveness observed under benzoate exposure.

From a bioprocess perspective, a progressive loss of cellular responsiveness under repeated or sustained benzoate exposure poses a direct threat to continuous culture stability. Unlike bulk variables such as dissolved oxygen or optical density, which reflect the consequences of collapse only after it has begun. Responsiveness, measured as the fraction of non-responsive cells in real time, may provide an early-warning signal that precedes irreversible biomass decline. Detecting and acting on this signal autonomously represents an opportunity to reframe bioprocess control around population structure rather than bulk physiology.

Here, we address three connected questions using *P. putida* KT2440 as a model system. First, does benzoate exposure reduce cellular responsiveness through a mechanism that is distinct from classical switching-cost-driven diversification, and does this effect scale with concentration and generalize across gene circuits? Second, can a resource allocation model (incorporating the competing demands of benzoate assimilation, toxicity, and tolerance) quantitatively explain the loss of responsiveness as an emergent consequence of limited cellular resources? Third, can real-time monitoring of cellular responsiveness via automated flow cytometry serve as an actionable control variable to prevent population collapse and enable autonomous continuous cultivation at otherwise destabilizing benzoate loads?

## Results

### 1. Different sources of cellular burden drive distinct diversification and responsiveness regimes in *P. putida*

To disentangle the effects of different sources of metabolic burden on population dynamics, we first established a reference regime using a synthetic rhamnose-inducible gene expression system as a tunable source of regulatory burden in *P. putida* KT2440, operated in the Segregostat platform (**Figure 1**). We previously developed the Segregostat to monitor cell population response based on the intrinsic dynamics of the biological system^7^ ^8^. When the rhamnose-inducible circuit was expressed without a growth decoupler, induction produced a homogeneous population-level response with homogenous fluorescence distributions over time and low population entropy, further confirming that low switching cost induces low entropy in *P. putida*.

**Figure 1:**
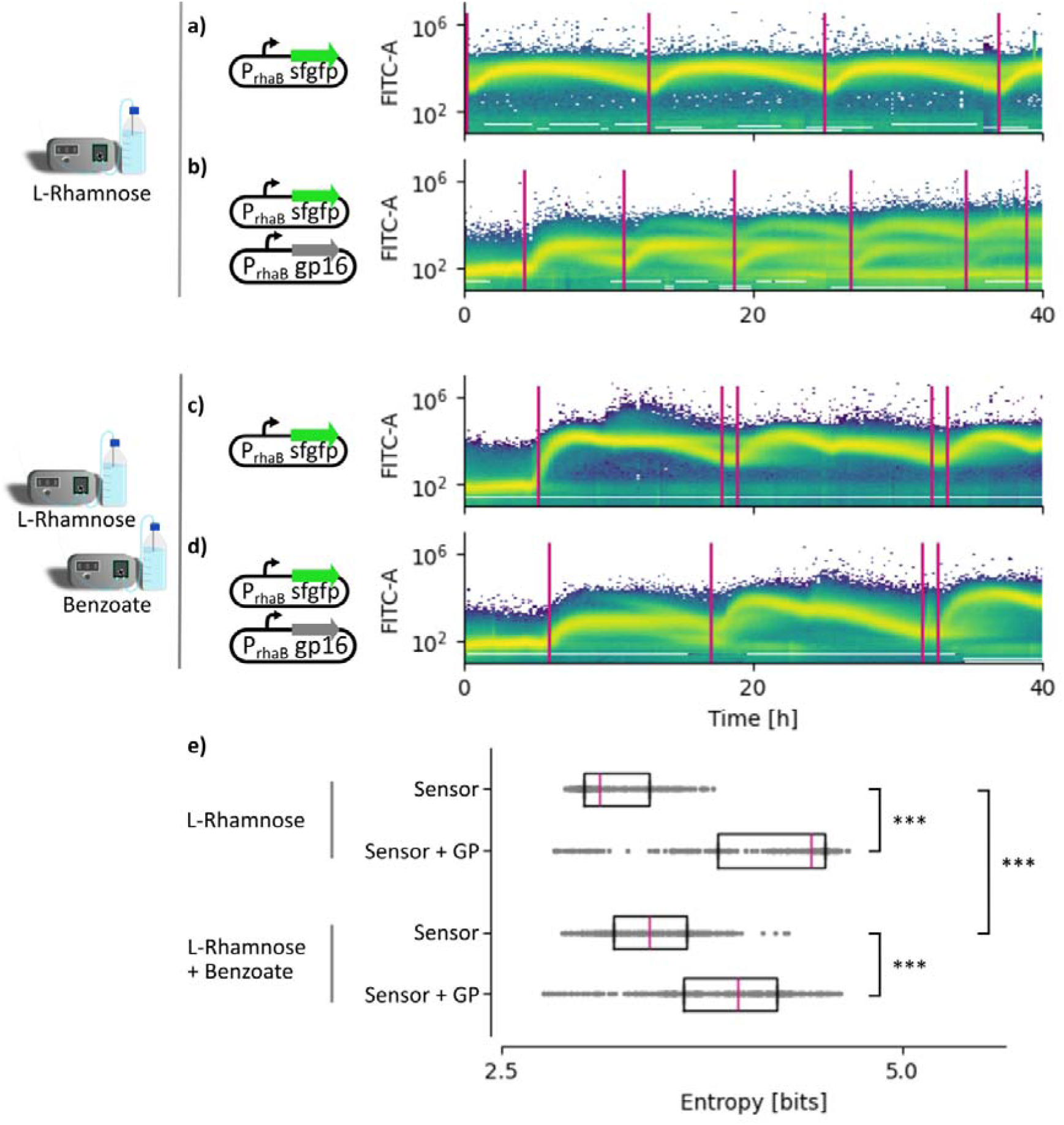
Switching cost related to the activation of burdensome gene circuits and benzoate exposure drive distinct population dynamics. **(A)** Population fluorescence distributions over time in *P. putida* KT2440 carrying a rhamnose-inducible reporter circuit expressed without a growth decoupler, operated in the Segregostat platform. Induction produces a homogeneous population response with low entropy, consistent with a low switching cost regime. **(B)** Introducing the growth-decoupling protein gp16 in parallel of the same circuit increases the fitness cost associated with reporter expression. Under these conditions, the population diversifies into a complex mixture of induced and non-induced cells, with population entropy markedly higher than control. **(C–D)** Addition of benzoate in Segregostat cultures at increasing concentrations does not promote phenotypic diversification. Population entropy remains unchanged or decreases relative to glucose-only controls, and fluorescence distributions do not resolve into multimodal patterns even at concentrations that reduce growth rate. **(E)** This absence of diversification persists even when benzoate is combined with co-expression of the growth-decoupler circuit, a condition that alone drives strong heterogeneity. Benzoate therefore operates through a mechanism distinct from classical switching-cost-driven diversification.

Introducing a growth decoupler on a second plasmid increased the fitness/switching cost associated with reporter expression^23^. Under these conditions, the population consistently diversified into a complex mix of induced and non-induced cells, with population entropy increasing markedly relative to the no-decoupler control (**Figure 1B**). This diversification pattern replicates and extends our previous findings and establishes a clear phenotypic benchmark i.e., high switching costs promote population heterogeneity as a buffering strategy, distributing the fitness penalty of induction across a spectrum of phenotypic states^7^.

We next asked whether an equivalent diversification response would arise when metabolic burden is imposed externally through a toxic substrate. Benzoate was added to glucose-grown Segregostat cultures at increasing concentrations while the rhamnose-inducible circuit was simultaneously induced. Contrary to expectations based on switching-cost rule, benzoate addition did not promote phenotypic diversification (**Figure 1C-D**). Population entropy remained unchanged or decreased relative to glucose-only controls, and the fluorescence distribution did not resolve into a multimodal pattern, even at benzoate concentrations that are supposed to reduce growth rate. Strikingly, this absence of diversification persisted even in conditions where co-expression of the growth-decoupler synthetic circuit alone would normally induce strong heterogeneity (**Figure 1E**). Benzoate therefore does not act according to the classical switching cost.

Analysis of the temporal population trajectories revealed that benzoate exposure affected the dynamics of circuit activation rather than its heterogeneity (**Figure 2**). Specifically, activation of the rhamnose-inducible reporter was delayed and attenuated following each induction event in the presence of benzoate. The maximum change in fluorescence i.e., our operational measure of cellular responsiveness, decreased progressively with increasing benzoate concentration. At low benzoate concentrations, the response was largely preserved. However, at high concentrations, circuit activation was delayed, with the population appearing transiently unable to engage the required physiological state despite the presence of inducer.

**Figure 2:**
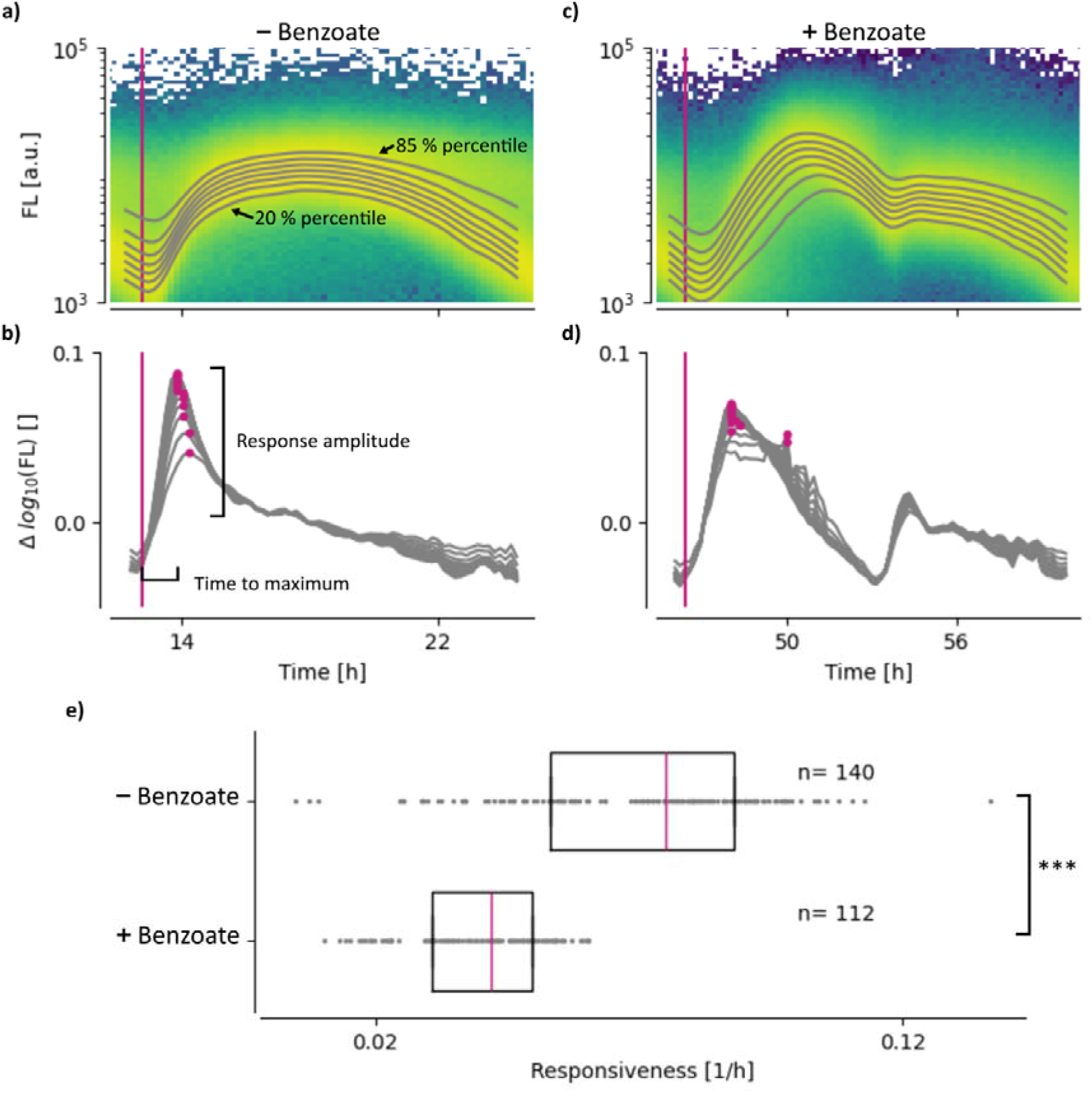
Addition of benzoate decreases cellular responsiveness. **(A–C)** Representative population trajectories as percentiles of fluorescence obtained from Segregostat experiments with *Pseudomonas putida* KT2440 carrying a rhamnose-inducible fluorescent reporter with **(A)** and without **(C)** benzoate. For each condition, the temporal evolution of the fluorescence distribution and the percentiles are shown following repeated induction events. (**B**) In the absence of benzoate, the change in fluorescence is fast, pointing out that gene circuit is quickly involved. (**D**) At high benzoate concentration, activation is markedly delayed and attenuated, with the population appearing transiently unable to engage the required physiological state despite the continued presence of inducer. **(E)** Quantification of cellular responsiveness for the two conditions, defined as the maximum change in fluorescence per induction cycle.

Together, these results demonstrate that not all forms of metabolic burden generate equivalent population responses. While fitness-costly gene expression promotes diversification for redistributing phenotypic states and buffering switching costs, benzoate suppresses the collective capacity to activate a gene circuit within a biologically relevant timescale following an environmental transition. This loss of responsiveness will be examined in the following section.

### 2. Loss of responsiveness scales with benzoate concentration and is correlated to fitness loss

The Segregostat experiments suggested that benzoate suppresses cellular responsiveness in a concentration-dependent manner. To determine whether this effect is substrate-specific and whether it is coupled to fitness loss, we performed microplate cultivations that enable simultaneous, high-throughput monitoring of reporter fluorescence and growth across a range of carbon sources and concentrations (**Figure 3A**). Four alternative carbon sources were tested alongside glucose i.e., succinate, ethylene glycol, acetate, and benzoate. Succinate and ethylene glycol served as non-inhibitory controls i.e., substrates that *P. putida* KT2440 assimilates efficiently without imposing measurable stress, while acetate and benzoate were selected as substrates associated with varying degrees of metabolic demand and potential toxicity.

**Figure 3:**
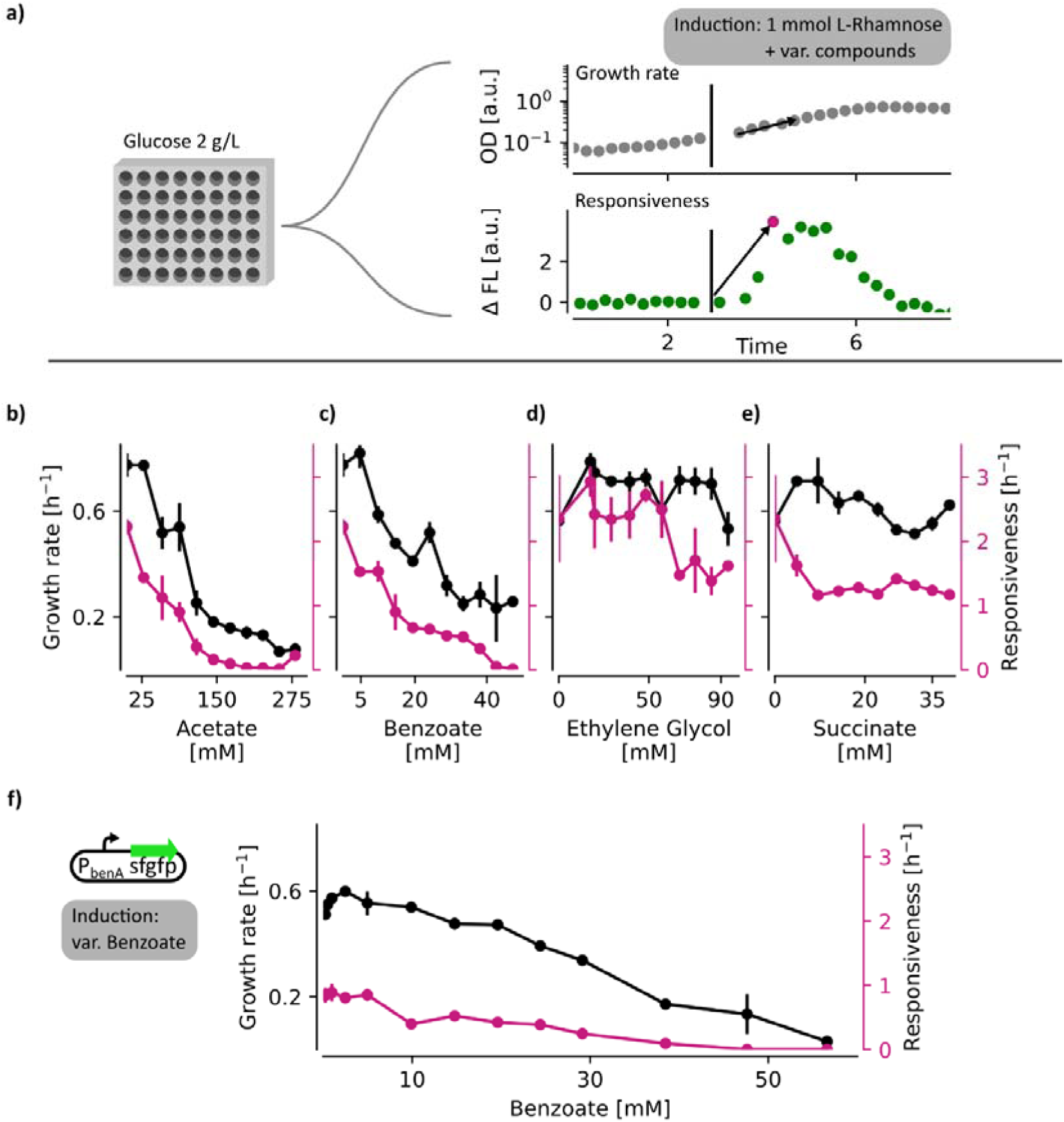
Loss of gene circuit responsiveness correlates with reduced cellular fitness. **(A)** Experimental design for high-throughput microplate cultivations monitoring reporter fluorescence and growth rate simultaneously across multiple carbon sources and concentrations. **(B–C)** Increasing concentrations of benzoate and acetate lead to a progressive and coupled decline in both cellular responsiveness and growth rate. The relationship is graded i.e., partial reductions in responsiveness are detectable before growth rate is affected, consistent with a resource competition mechanism. **(D–E)** Succinate and ethylene glycol, used as non-inhibitory controls, do not affect cellular responsiveness or growth rate relative to glucose across the concentration range tested. **(F)** Responsiveness of the native benzoate-responsive *benA* reporter shows an equivalent concentration-dependent impairment upon benzoate exposure, demonstrating that the loss of responsiveness is not specific to the regulatory architecture of the heterologous rhamnose-inducible system but reflects a host-level physiological constraint that limits transcriptional activation across both native and synthetic gene circuits.

For succinate and ethylene glycol, neither cellular responsiveness nor growth rate was affected relative to glucose controls across the concentration range tested (**Figure 3D-E**). By contrast, increasing concentrations of both acetate and benzoate led to a pronounced and progressive decrease in cellular responsiveness, concomitant with a significant reduction in growth rate (**Figure 3B-C**). Under these conditions, both responsiveness and growth rate declined in parallel across the concentration range. Importantly, the relationship was graded rather than threshold-like i.e., even at sub-inhibitory concentrations, partial reductions in responsiveness were detectable before growth rate was substantially affected. This observation is consistent with the resource allocation hypothesis that will be developed in the following section and suggesting that responsiveness is lost as resources are progressively diverted toward substrate assimilation and stress mitigation before the growth rate itself collapses.

To assess whether the loss of responsiveness was an artefact of the heterologous rhamnose-inducible circuit, we quantified the response of the native benzoate-responsive operon under equivalent conditions. Using a *benA* reporter strain in which the endogenous promoter drives fluorescent protein expression, we observed a similar concentration-dependent loss of responsiveness upon benzoate exposure (**Figure 3F**). Segregostat experiments performed with the *benA* reporter at 1 and 20 mM benzoate further confirmed this behavior (**Supplementary Information, Figure S2**). At low concentrations, the native circuit activated rapidly and completely following substrate addition whereas, at high concentrations, activation was delayed, and in some cases absent within the observation window. This result demonstrates that the impairment of responsiveness is not specific to the regulatory architecture of the heterologous system but reflects a host-level physiological constraint that limits transcriptional responsiveness across both heterologous and native gene circuits.

### 3. A resource allocation model explains responsiveness loss as a competition for limited cellular resources

The previous sections established that benzoate reduces cellular responsiveness in a concentration-dependent manner correlated with fitness loss. However, the experiments do not directly reveal why benzoate impairs responsiveness. To address this mechanistic question, we developed a resource allocation model that connects the observable population behaviour i.e., growth rate, substrate consumption, and circuit activation, to intracellular allocation decisions that cannot be measured directly in our experimental system.

Resource allocation models describe cellular phenotypes as emergent properties of how a finite pool of biosynthetic capacity is distributed among competing functional sectors ^1^ ^2^. In *E. coli*, such models have shown that suboptimal allocation during environmental transitions directly constrains both growth recovery and gene circuit activation, providing experimental evidence that resource competition, rather than specific regulatory wiring, can be the primary bottleneck during substrate shifts ^2^. Extensions of this framework have since captured how antibiotic stress forces a trade-off between growth and tolerance ^20^, and how resource partitioning shapes heterologous gene expression in *P. putida* under nutrient limitation ^22^ ^21^. Our model builds on this foundation but addresses a new scenario by considering a substrate that simultaneously demands investment in a dedicated assimilation sector, imposes concentration-dependent toxicity, and requires the formation of a tolerance sector.

The model represents cellular biomass as partitioned among four functional sectors (**Figure 4A**). Substrate uptake, from a non-inhibitory carbon source (S, glucose) and an inhibitory substrate (B, benzoate), generates an unspecialized resource pool (R) that serves as the currency for all downstream allocation. This unspecialized biomass is then distributed, at an allocation cost, among three specialized sectors i.e., a metabolic sector for glucose assimilation (Cs), a metabolic sector for benzoate assimilation (Cb), and a tolerance sector (T) that counteracts the inhibitory effects of benzoate on cellular viability. Total biomass is the sum of these four compartments, and each metabolic sector regulates the uptake rate of its corresponding substrate. Importantly, allocation toward any specialized sector reduces the resources available to all others, so that investment in benzoate assimilation or tolerance necessarily competes with the capacity to maintain or expand the adaptive machinery required for responsiveness.

**Figure 4:**
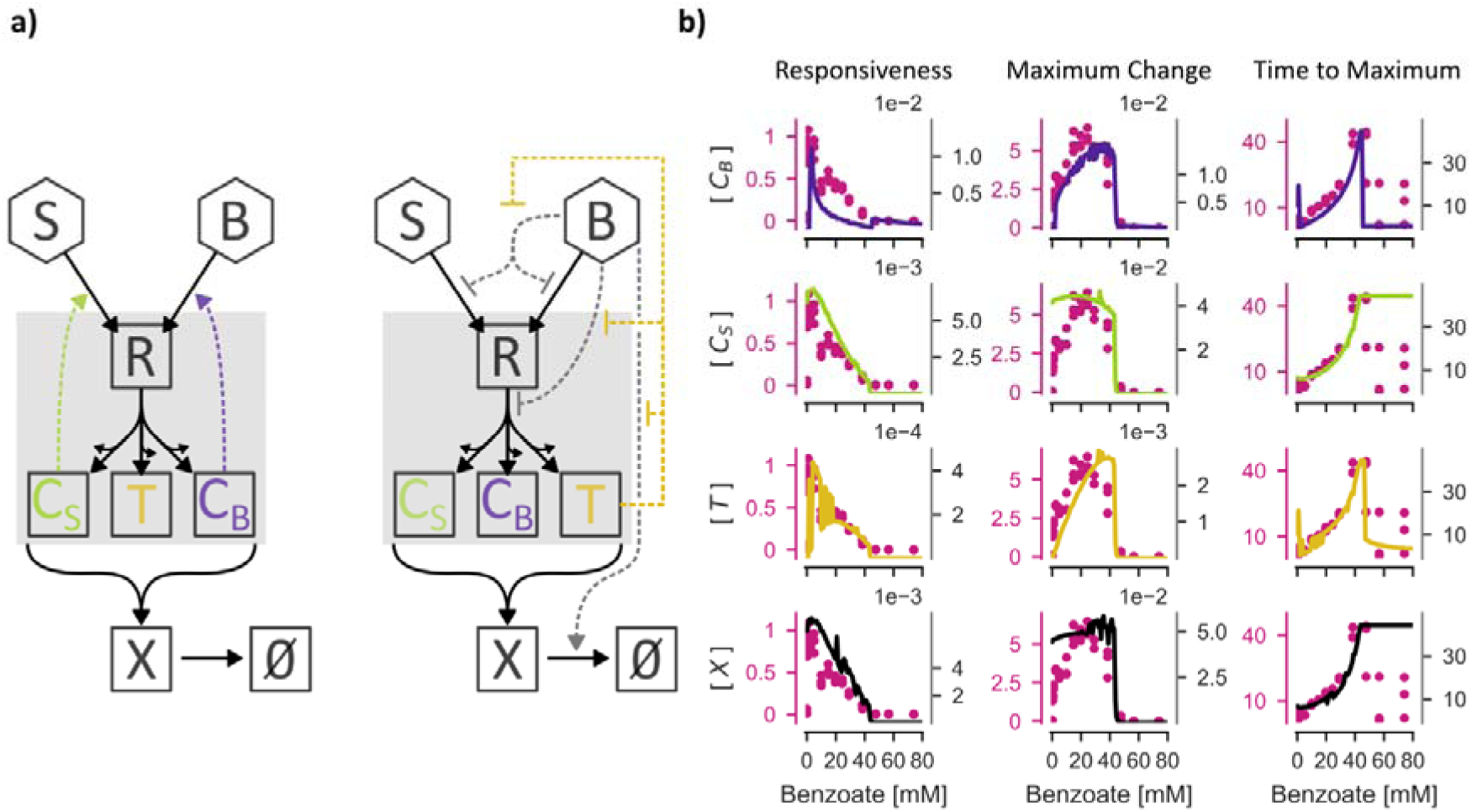
A resource allocation model explains loss of responsiveness as an emergent consequence of competition for limited cellular resources. **(A)** Schematic of the four-sector resource allocation model. Cellular biomass is partitioned among an unspecialized resource pool (R) generated by substrate uptake from glucose (S*g*) and benzoate (S*b*), and three specialized sectors i.e., a metabolic sector for glucose assimilation (C*g*), a metabolic sector for benzoate assimilation (C*b*), and a tolerance sector (Z) that counteracts benzoate-induced inhibition. Allocation toward any specialized sector reduces resources available to all others, so that investment in benzoate assimilation and tolerance directly competes with the adaptive reallocation capacity required for circuit activation. **(B)** Model simulations predict a critical benzoate concentration above which the three-way competition between assimilation, tolerance, and adaptive reallocation exhausts the unspecialized resource pool, causing responsiveness to collapse. Below this threshold, sufficient resources remain for both tolerance formation and circuit activation. Simulated biomass trajectories are overlaid with experimental batch culture data collected across a range of benzoate concentrations, showing accurate reproduction of both the progressive reduction in final biomass yield and the extended lag phase observed at high concentrations.

Benzoate inhibition is incorporated at three levels. First, it reduces substrate uptake rates in a concentration-dependent manner, capturing the growth inhibition observed experimentally. Second, it reduces the rate of allocation toward specialized sectors, reflecting the impairment of resource redistribution that we hypothesize underlies the loss of responsiveness. Third, at high concentrations, benzoate increases protein denaturation and degradation rates, directly impacting cellular viability. The tolerance sector counteracts these effects since its formation stabilizes growth under benzoate exposure by neutralizing inhibitory damage. However, building the tolerance sector requires resources drawn from the same unspecialized pool, creating a direct competition between tolerance and the adaptive reallocation required for circuit activation.

Model simulations point out a critical benzoate concentration above which cellular responsiveness is lost due to this three-way resource competition (**Figure 4B**). Below this threshold, the tolerance sector can be formed rapidly enough to stabilize growth while leaving sufficient resources for adaptive reallocation, so that responsiveness is preserved even as growth rate declines. Above the threshold, the demand imposed by simultaneous assimilation and tolerance formation exhausts the unspecialized resource pool, leaving the cell unable to reallocate resources toward the activation of new physiological states within a biologically relevant timescale. Responsiveness therefore collapses as a result of these allocations issues. This prediction is consistent with the graded, concentration-dependent loss of responsiveness observed experimentally in both the microplate and Segregostat assays (**Figure 3**). To validate this prediction, we compared model simulations against batch culture data collected across a range of benzoate concentrations. The model accurately reproduced the experimentally observed biomass trajectories, capturing both the progressive reduction in final biomass yield and the extended lag phase at high benzoate concentrations (**Figure 4B**).

### 4. Repeated environmental transitions with benzoate drive population collapse in continuous culture

Our results point out that benzoate reduces cellular responsiveness and fitness in a concentration-dependent manner, while ethylene glycol (a substrate of comparable biotechnological interest as a plastic-derived feedstock) leaves both variables largely unaffected. We therefore used these two substrates to investigate whether the responsiveness–fitness coupling observed under static conditions would manifest as a difference in population stability under continuous cultivation. Experiments were designed to impose repeated environmental transitions by pulsing either benzoate (**Figure 5A&C**) or ethylene glycol (**Figure 5B&D**) into glucose-limited continuous cultures, at increasing pulsing frequencies (**Supplementary Information, Table S6**). At low pulsing frequencies, both substrates elicited sustained oscillatory dynamics in population fluorescence and biomass, indicating that cells were capable of repeatedly engaging and disengaging the metabolic program required for assimilation of the alternative carbon source (**Figure 5**). This oscillatory behaviour was reproducible across multiple pulsing cycles and was not associated with any measurable decline in biomass or dissolved oxygen, confirming that low-frequency transitions are well tolerated regardless of the substrate. Increasing the pulsing frequency led to a progressive damping of the oscillatory response in both cultures. However, the two substrates diverged markedly in the consequences of this damping. For ethylene glycol, attenuation of the oscillatory amplitude remained largely neutral with respect to biomass and substrate consumption i.e., the population adapted to higher transition frequencies without a measurable decline in overall fitness, and dissolved oxygen remained stable throughout the cultivation (**Figure 5D**). For benzoate, damping was accompanied by a progressive and irreversible loss of cellular responsiveness that translated into biomass decline and failure of substrate assimilation (**Figure 5C**). The rise in dissolved oxygen observed toward the end of benzoate cultivations at high pulsing frequency provides unambiguous evidence that the culture was no longer consuming benzoate at a rate consistent with sustained growth.

**Figure 5:**
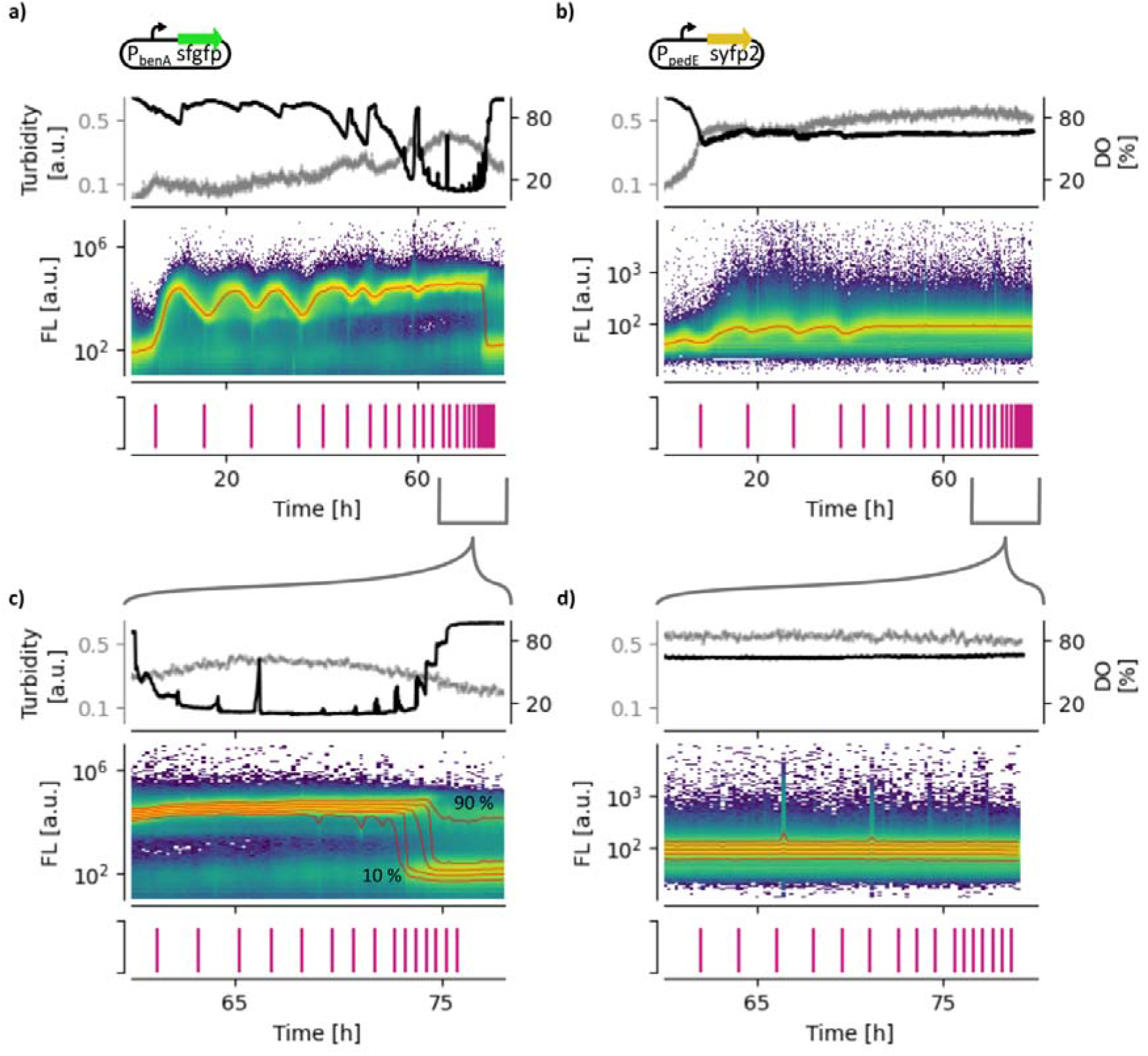
Substrate-dependent collapse of cellular responsiveness in continuous cultures. **(A & C)** Continuous cultivation of *P. putida* KT2440 under repeated benzoate pulsing at increasing frequencies. At low pulsing frequency, sustained oscillatory dynamics in population fluorescence are observed, indicating repeated engagement and disengagement of the benzoate assimilation program. At high pulsing frequency, oscillatory amplitude is progressively damped and accompanied by an irreversible loss of cellular responsiveness. At this stage, biomass decline and substrate assimilation fails, as evidenced by a rise in dissolved oxygen. Percentile-based analysis of flow cytometry data resolves a responsive fraction i.e., cells that continue to activate the biosensor reporter following each pulse and a non-responsive fraction i.e., cells that fail to cross the activation threshold within the observation window. The non-responsive fraction accumulates consistently before the onset of macroscopic collapse, preceding any detectable rise in dissolved oxygen or decline in optical density, establishing it as an early-warning indicator of culture instability. **(B & D)** Equivalent continuous cultivation under repeated ethylene glycol pulsing at increasing frequencies. Oscillatory dynamics are similarly damped at high pulsing frequency, but attenuation remains neutral with respect to biomass and substrate consumption. The population adapts to higher transition frequencies without measurable fitness decline, and dissolved oxygen remains stable throughout, demonstrating that the collapse phenotype observed under benzoate is substrate-specific and not a generic consequence of increased transition frequency.

A notable feature of the collapse dynamics under benzoate pulsing was the consistent emergence of two phenotypically distinct subpopulations in the period preceding and following complete biomass decline (**Figure 5C**). Percentile-based analysis of the flow cytometry data resolved a responsive fraction i.e., cells that continued to activate the biosensor reporter following each benzoate pulse, and a non-responsive fraction i.e., cells that failed to cross the activation threshold within the observation window. The non-responsive fraction began accumulating before the onset of collapse as measured by dissolved oxygen or optical density.

This pattern of collapse is fully consistent with the resource allocation model developed in the previous section. The model predicted that above a critical benzoate concentration, cells lose the allocation capacity to engage new physiological states even though they retain the metabolic potential to tolerate higher loads (**Figure 4**). In continuous culture under repeated pulsing, each successive transition incrementally depletes the resource available for adaptive reallocation, progressively shifting cells from the responsive to the non-responsive phenotypic state. Once the non-responsive fraction reaches a critical threshold, the population can no longer sustain benzoate assimilation at the rate required for balanced growth, and collapse is observed.

### 5. Real-time monitoring of subpopulation structure enables autonomous prevention of process collapse

The continuous culture experiments established that the emergence of a non-responsive subpopulation is an early-warning signal of collapse under high benzoate loads. This raises the question whether this signal can be exploited to prevent collapse while maintaining effective benzoate assimilation? A control strategy based on bulk variables such as dissolved oxygen or optical density would be reactive, intervening only after collapse has begun. By contrast, a strategy based on population structure, specifically the fraction of non-responsive cells, would be preventive and would detect the precursor signature and adjust substrate feeding before irreversible damage to the culture occurs. To test this concept, we designed a two-stage connected bioreactor system that couples real-time population monitoring to autonomous feed control.

The system consisted of two bioreactors connected in series (**Figure 6A**). The first vessel was operated under Segregostat control i.e., automated flow cytometry measurements were taken at regular intervals, and benzoate pulsing decisions were made in real time based on the current population structure. Specifically, benzoate addition was allowed if the fraction of non-responsive cells in the first vessel remained below a predefined threshold of 10% (**Figure 6B&C**). When this threshold was exceeded, indicating that the population was approaching the collapse regime, benzoate pulsing was automatically stopped, allowing the population to recover responsiveness before the next addition. The outlet stream of the first vessel was directed into a second vessel operating under standard continuous cultivation conditions, receiving a baseline feed of glucose and low-concentration benzoate.

**Figure 6:**
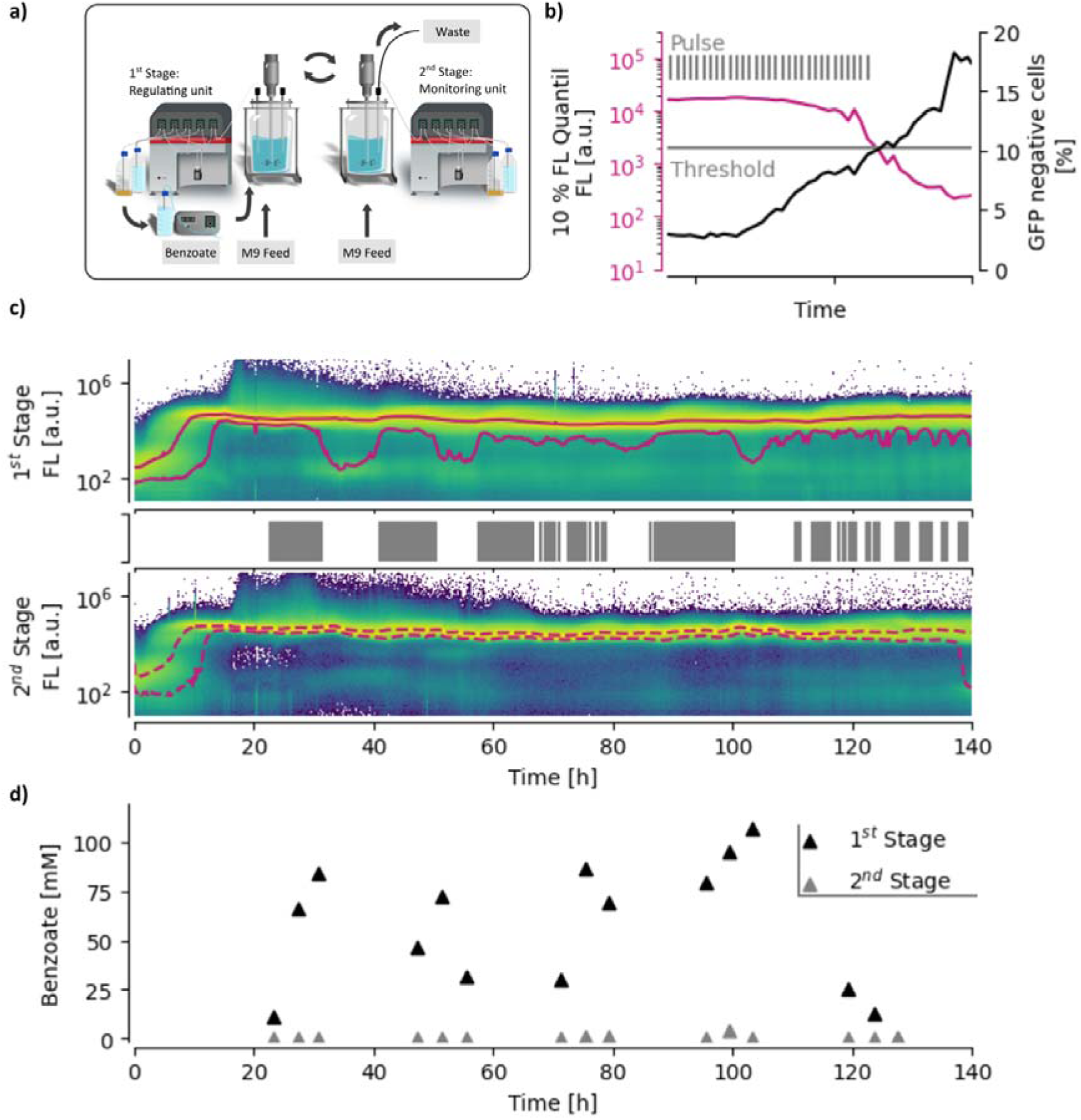
Cellular responsiveness-based control enables autonomous cultivation at high benzoate concentrations. **(A)** Schematic of the two-stage connected bioreactor system. The first vessel is operated under Segregostat control with automated flow cytometry measurements taken at regular intervals. The second vessel operates under standard continuous cultivation conditions, receiving a baseline feed of glucose and low-concentration benzoate. The outlet stream of the first vessel is directed into the second vessel, and the return flow of cells provides a source of responsive phenotypes that replenishes the first vessel during recovery periods. **(B–C)** Real-time population structure monitoring in the first vessel. Benzoate addition is allowed when the fraction of non-responsive cells remains below 10%. When this threshold is exceeded, pulsing is automatically halted to allow population recovery before the next addition cycle. **(D)** Benzoate concentration profiles across the cultivation period demonstrating complete substrate consumption under population-structure-based control, contrasting with the accumulation of residual benzoate and biomass collapse observed under uncontrolled high-benzoate conditions. Together, these results establish cellular responsiveness as a quantitative, actionable control variable enabling stable autonomous bioprocess operation on a challenging aromatic feedstock.

This population-structure-based control strategy achieved two outcomes that could not be obtained simultaneously under uncontrolled high-benzoate conditions. First, it enabled complete benzoate consumption across the cultivation period (**Figure 6D**) by preventing the accumulation of a non-responsive majority. This strategy ensured that a sufficient responsive fraction was always present to assimilate each benzoate pulse before the next measurement cycle. Second, it prevented population collapse in the first vessel despite repeated exposure to benzoate concentrations that caused irreversible biomass decline in non-controlled cultivation. Additionally, the return flow of cells from the second vessel to the first vessel (via the shared medium circuit) effectively functions as a source of responsive phenotypes that replenishes the first vessel during recovery periods. Together, these results demonstrate that cellular responsiveness i.e., a population-level property that is invisible to bulk process sensors, can be monitored in real time and used as an actionable control variable to sustain autonomous continuous cultivation on a challenging substrate.

## Discussion

A wide range of cellular constraints are known to generate fitness trade-offs that shape gene expression capacity and population structure. Ribosome limitation, metabolic burden imposed by synthetic gene circuits, and resource competition between heterologous and endogenous processes all reduce the resources available for growth and can promote phenotypic diversification, a strategy in which fitness costs are distributed across a spectrum of coexisting phenotypic states ^7^ ^24^ ^2^ ^19^. In such systems, transitions between phenotypic states carry measurable switching costs, and populations evolve toward heterogeneity ^7^ ^25^. The present work reveals that not all sources of fitness trade-offs generate this diversification response. Benzoate exposure under dynamic conditions operates on a different axis. Rather than increasing population entropy, it progressively reduces cellular responsiveness i.e., the capacity of the population to activate the appropriate metabolic program following an environmental transition. These two outcomes, diversification and loss of responsiveness, are mechanistically separable and substrate-specific, and they lead to qualitatively different population dynamics in continuous culture. Recognizing this distinction is important both for understanding microbial physiology under complex substrate regimes and for designing bioprocesses that operate reliably on challenging feedstocks.

The resource allocation model developed here provides a mechanistic framework that connects these observations. By treating the cell as a system that partitions a finite biosynthetic capacity among competing functional sectors i.e., substrate assimilation, tolerance, and adaptive reallocation, the model point out that loss of responsiveness is a predictable consequence of resource competition rather than a benzoate-specific regulatory effect. This framing extends prior resource allocation models that have explained growth-expression trade-offs in *E. coli* ^2^ and antibiotic tolerance in other systems^20^, by incorporating the three-way competition created when a substrate simultaneously demands assimilation machinery, imposes toxicity, and requires a dedicated tolerance sector. The model predicts a critical benzoate concentration above which responsiveness collapses. Interestingly, this threshold is detectable at the population level via biosensor-based flow cytometry. This illustrates the fact that resource allocation models can predict qualitative transitions in population behaviour that are invisible to bulk physiological measurements and can therefore serve as a design tool for identifying operating regimes in which population stability is at risk.

A key observation of this study is the consistent emergence of a phenotypically distinct non-responsive subpopulation in the period preceding population collapse under high-frequency benzoate pulsing. The mechanism behind the formation of this subpopulation can have different sources. One interpretation, consistent with the resource allocation model, is that non-responsive cells are not genetically distinct but represent a physiological state in which resource depletion has rendered adaptive reallocation temporarily impossible i.e., cells that have invested heavily in tolerance under repeated benzoate exposure are transiently unable to redirect resources toward circuit activation and remain locked in a non-responsive state until resource availability is restored. A second interpretation is that non-responsive cells actively downregulate benzoate uptake as a stress-avoidance strategy, effectively shutting down substrate import to limit intracellular toxicity at the cost of reduced assimilation capacity.

Regardless of its mechanistic origin, the non-responsive subpopulation consistently emerged before macroscopic collapse before the rise in dissolved oxygen and before measurable decline in biomass, positioning it as a reliable early-warning indicator of instability. This temporal ordering is the observation that makes population-structure-based control both possible and advantageous. Conventional process sensors e.g., dissolved oxygen, optical density, pH, detect the failure after it has begun. The non-responsive fraction, by contrast, reflects the accumulation of switching costs and the progressive decline of adaptive capacity preceding collapse. Monitoring it therefore converts a reactive control problem into a proactive one, enabling intervention at a point where the population retains sufficient responsive cells to recover. More broadly, this result supports the emerging view that single-cell resolved population metrics, accessible through real-time flow cytometry and biosensor-equipped strains, can serve as early-warning indicators in bioprocesses, bridging the gap between molecular-level physiology and process-level stability ^26^ ^27^ ^28^ ^29^.

The control strategy implemented here exploits this early-warning signal to sustain autonomous continuous cultivation under high benzoate loads that would otherwise cause collapse. By coupling automated FC monitoring with a dynamic benzoate feeding policy, we achieved complete substrate consumption and stable population dynamics over extended cultivation times. Importantly, this strategy does not attempt to suppress phenotypic heterogeneity and accepts that a non-responsive subpopulation will emerge under high benzoate load and uses its abundance as the control trigger. This aligns with an emerging paradigm in cybergenetics and feedback-controlled bioprocesses, in which phenotypic state distributions, rather than bulk variables alone, guide autonomous regulation ^30^ ^31^ ^32^ ^33^. The present work extends this paradigm to a scenario where the relevant phenotypic distribution is not engineered but emergent, arising from the interplay between substrate uptake physiology, resource allocation, and population dynamics and demonstrates that emergent heterogeneity can be harnessed as a stabilizing control handle rather than treated as noise to be minimized.

Taken together, these observations position structure-aware, population-resolved control as an interface between microbial physiology, systems and synthetic biology, and bioprocess engineering.

## Material and methods

### Strains and plasmids

For this study, the model strain *Pseudomonas putida* KT2440 was used and equipped with specific biosensors (**Supplementary Information, Table S1**). The biosensors were based on the pSEVA231 plasmid ^34^ carrying a kanamycin resistance marker. The vector was first linearized using the restriction enzyme *Bam*HI, and the fluorophore sequence *sfgfp-rrnB* was inserted by HiFi assembly. The resulting plasmid, pSEVA231·sfgfp, contains a *Bam*HI restriction site upstream of the fluorophore. This site was subsequently used to introduce specific expression cassettes by HiFi assembly (RhaRS-P_RhaBAD_, P_benA_, and P_pedE_, **Supplementary Information, Table S2**). These constructs allow monitoring of the native expression associated with the utilization of benzoate and ethylene glycol (EG) ^35^ ^36^ ^37^ ^17^, as well as the activity of a heterologous gene circuit not present in *P. putida* KT2440 ^38^ ^39^. The heterologous system is based on L-rhamnose induction. For the EG biosensor, the fluorophore *sfgfp* was replaced with *syfp2*. The plasmid pSEVA231·PpedE-sfgfp was linearized by PCR using the primer pair pSEVA231·sfgfp-Backbone-F/R and assembled with the *syfp2* fragment. The promoter fragments for P_benA_ and P_pedE_ were synthesized by Integrated DNA Technologies. The expression cassette RhaRS-P_RhaBAD_ was amplified using the primer pair Insert-P_RhaBAD_-F/R from the plasmid pSTDesR·phi15gp16. This plasmid was originally constructed for the expression of the growth-decoupling protein gp16, a phage-derived protein that inhibits the host RNA polymerase ^23^. In this study, gp16 expression was used as a synthetic burden controlled by L-rhamnose induction.

### Plasmid construction and transformation

The plasmid amplification and transformation procedure was adapted from Wirth *et al.* ^40^. Plasmids were amplified in *E. coli* DH5α in 100 mL baffled shake flasks containing 10 mL of LB medium supplemented with the appropriate antibiotics (50 ng/µL kanamycin or 100 ng/µL streptomycin) at 37 °C. Plasmid assembly was performed using the NEBuilder^®^ HiFi DNA Assembly Master Mix from New England Biolabs, with 20 bp overhangs on both sides. For the electroporation, an overnight culture of *P. putida* was aliquoted into 1 mL portions in 2 mL reaction tubes and centrifuged for 1 min at 11,000 *g*. The cell pellets were resuspended in 1 mL of 300 mM sucrose solution. This washing step was repeated twice. After the final wash, the pellets were resuspended in 100 µL of 300 mM sucrose solution. Electroporation was performed using 2 µL of plasmid assembly mixture and 100 µL of competent cells at 25 µF, 2.5 kV, and 200 Ω in cuvettes with a 2 mm gap. Cells were immediately resuspended in 1 mL of LB medium and incubated at 30 °C for 2 h. Finally, 100 µL of the culture was plated on LB agar plates containing the appropriate antibiotics and incubated overnight at 30 °C. The remaining 900 µL were concentrated by centrifugation and incubated identically.

### Culture conditions and media

Cells were cultured either in LB or in minimal salt media. The minimal salt media contained 14.6 g/L K_2_HPO_4_, 3.6 g/L NaH_2_PO_4_·2 H_2_O, 2 g/L (NH_4_)_2_SO_4_, 2.47 g/L NH_4_Cl, 1 g/L NH_4_Cl and 11 mL of a sterile-filtered trace elements solution. The trace elements solution contained within 1 L: 6.03 g EDTA, 5.01 g FeCl_3_·6 H_2_O, 36 g MgSO_4_, 1 g thiamine hydrochloride, 222 mg CaCl·2 H_2_O, 54 mg ZnSO_4_ ·7 H_2_O, 30 mg MnSO_4_·H_2_O, 30 mg CuSO_4_·5 H_2_O, and 63 mg CoSO_4_·7 H_2_O. Further, the media were supplemented with the appropriate antibiotic (50 ng/µL kanamycin or 100 ng/µL spectinomycin; stock solutions at 50 µg/µL and 100 µg/µL, respectively). Glucose or other carbon sources were added as described in the text.

For the preculture, cells were first cultured on LB plates overnight at 30 °C. One colony was used to inoculate 10 mL of LB medium in 100 mL baffled shake flasks. After 6 h, 500 µL was transferred to 50 mL of minimal salt medium (10% working volume, baffled flask) containing 10 g/L glucose. Experiments started on the following day with an initial OD_600_ of 0.1. Characterization in multiplate readers was performed with Spark® Multimode Microplate Reader (Tecan) using either 48 Well (BioLite 48 Well Multidish, Thermo Scientific) or 96 Well plates (TC-Platte 96 Well, Standard F, Sarstedt AG & Co.KG) at a working volume up to 0.5 mL or 0.2 mL, respectively. The temperature was maintained at 30 °C. Each measurement cycle included Absorbance at 600 nm (OD_600_) and fluorescence measurements (bandpass filter Ex: 485/20, Em: 535/35), followed by 900 s of shaking. The fluorescence signal was normalized by dividing by the mean fluorescence of a culture containing the biosensor without an inducer.

Bioreactor cultivations were performed in the F1 lab-scale bioreactor (Bionet) at 30 °C, pH = 7, an aeration rate of 1 L/min, a stirrer speed of 1000 rpm, and 1 g/L glucose. Turbidity was measured online by the Dencytee Arc RS485 probe (Hamilton). When glucose was depleted (∼5 hours), the continuous process was initialised using the same minimal salt medium, with 1 g/L glucose as the feed, at a dilution rate of 0.3 h^-1^. Samples were centrifuged for 1 min at 2000 *g*, and the supernatant was stored at -20 °C.

Flow cytometer measurements were conducted by an autonomous sampling unit. The operation can be monitoring or combined with a regulatory operation. Based on the population distribution, an actuator action can be defined. This cell-machine interface, called Segregostat, has been described previously ^8^ ^9^. The sampling unit takes a sample every 10 to 15 min and dilutes it accordingly. The flow cytometer (BD Accuri C6+, BD Biosciences) measures cells with FSC-H > 15000 a.u. and GFP and YFP in the FL1 channel using a bandpass filter of 533/30 or 510/15, respectively. The feedback regulation based on the FC data is employed by a custom MATLAB script.

### Connected bioreactor experiment

For the reconnected cascade bioreactor, the modified salt medium was supplemented with 1 g/L (NH_4_)_2_-H-citrate; the carbon source in the batch was 30 mM benzoate, and in the feeding medium were 1 mM benzoate and 1 g/L glucose. The reactor configuration contained two reactors, with individual media inputs at flow rates of 30 mL/h and 60 mL/h, respectively. 5 mmol of benzoate (1 M stock solution, 4-5 pulses per hour) was delivered to the 1^st^ stage if no collapse was detected. A collapse was detected when more than 10% of the cells had fluorescence below 2000 a.u. The reactor volume was maintained at 1 L by an overflow standpipe connected to a pump at a flow rate of 250 mL/h, enabling reactive efflux in response to varying inflow rates. (**Supplementary Information, Figure S5**).

### Determination of benzoate concentration

Bioreactor supernatant was analyzed using a Shimadzu Nexera UPLC system equipped with an SCL-40 controller and a diode array detector (DAD). Separation was performed on a Waters ACQUITY UPLC BEH C18 column (1.7 µm, 2.1 × 50 mm) maintained at 40 °C. Samples were filtered through a 0.22 µm pore-size filter prior to injection. The mobile phase was delivered at a flow rate of 0.6 mL min⁻¹ and consisted of (A) water containing 0.1% trifluoroacetic acid and (B) acetonitrile containing 0.1% trifluoroacetic acid. The total run time was 17 min and consisted of the following steps: (i) injection of 10 µL sample and equilibration for 1.5 min at 100% A; (ii) gradient elution to 50% B over 10 min; (iii) column washing by increasing B to 100% within 2 min and maintaining this condition for 2 min; and (iv) column re-equilibration by decreasing B to 0% within 0.3 min and maintaining this condition until 17 min. Benzoate was detected at 214 nm.

### Calculation of Responsiveness

Responsiveness was used as a quantitative measure of the ability of a microbial population to react to an environmental perturbation. In this study, the perturbation consisted of the addition of an inducer (L-rhamnose or benzoate), resulting in the activation of GFP expression. Responsiveness was defined as the maximum rate of change in fluorescence intensity relative to the time required to reach this maximum response. Accordingly, responsiveness (Res) was calculated as:

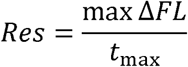

Where (*FL*) denotes fluorescence intensity and (*t_max_*) is the time at which the maximum fluorescence change is observed.

For Segregostat cultivations, flow cytometry fluorescence distributions were acquired at discrete sampling intervals. To characterize population heterogeneity, fluorescence percentiles were extracted from each distribution and transformed using the logarithmic scale. The resulting percentile trajectories were smoothed using a moving average window of five samples. Responsiveness was subsequently calculated for each percentile independently, allowing the determination of the population-level dispersion in dynamic responses.

For plate reader experiments, fluorescence measurements were first blank-corrected and normalised to the corresponding non-induced control condition. The change in normalised fluorescence was then calculated and used to determine responsiveness according to the procedure above.

### Resource allocation model

To describe cellular adaptation to benzoate stress, we developed a resource allocation model that captures growth on glucose and benzoate while accounting for substrate toxicity and tolerance development. The model partitions biomass into four functional sectors: an unspecified resource pool (*R*), glucose-utilizing biomass (*C_S_*), benzoate-utilizing biomass (*C_B_*), and a tolerance sector (*T*). Resource allocation between these sectors determines substrate uptake capacity, growth, and stress resilience. Benzoate inhibition is modelled as an uncompetitive process that is mitigated by investment in the tolerance sector. A complete model description is provided in **Supplementary Note S1**.

Model parameters were estimated using 45 growth curves obtained under 15 benzoate pulse concentrations (**Supplementary Fig. S3i**). Parameters were fitted using a two-stage Differential Evolution optimization procedure in SciPy. The estimation was performed on 80% of the dataset, while the randomly selected remaining 20% was used for validation. The objective function minimized the mean squared percentage error between model predictions and experimental observations.

## Acknowledgement

FD and FF received funding from the Action de Recherche Concertée (ARC) from ULiège (Plastinsect). MS received a PhD grant from the same ARC project. FD and LJ were supported by the Walloon Region and the Fond Européen de Développement Régional (FEDER) [Grant portfolio PHENIX n°244, project PHENIX_Technologique_ULiège n°397] and [Grant portfolio PHENIX n°244, project PHENIX_FoodBooster_ULiège n°428]. FD gratefully acknowledge the support and funding from the FRS-FNRS (project no. 40013556, WEAVE collaboration network). The authors also very gratefully to Prof. Rob Lavigne (KULeuven, Belgium) for the donation of the rhamnose-inducible systems and the growth decoupler.

## Notes

### Competing Interest Statement

The authors have declared no competing interest.

