## Supplementary material for "Cellular responsiveness as a predictive indicator for population collapse and autonomous control in continuous cultures of *Pseudomonas putida*"

**Supplementary Table S1. Strain used in this work.**

| **Name** | **Description** | **Reference** |
| --- | --- | --- |
| *P. putida* KT2440 | Derivative of P. putida mt-2 lacking the TOL plasmid | (Bagdasarian et al., 1981) |
| *E. coli DH5α* | Intermediate host for vector cloning  *fhuA2::IS2 Δ(mmuP-mhpD)169 ΔphoA8 glnX44 ϕ80d[ΔlacZ58(M15)] rfbD1 gyrA96 luxS11 recA1 endA1 rphWT thiE1 hsdR17* |  |

**Supplementary Table S2. Sequences and primers used in this work.**

| **Name** | **Sequence (5’-3’)** |
| --- | --- |
| SEVA_Seq-F | GTTGTCCACGGGCCGAG |
| SEVA_Seq-R | CGTAATTATTGGGGACCCCTGG |
| pSEVA231·*sfgfp-*Backbone-F | GCGGTTCTGAATTCACACCTAGGTAAC |
| pSEVA231·*sfgfp-*Backbone-R | GGGTTGCAGTTCCCAGTG |
| Insert-P_RhaBAD_-F | TTCGAGCTCGGTACCCGGGGggctctgctgtagtgagtgg |
| Insert-P_RhaBAD_-R | gttcttcgcccttgctcatgtgtcgaccacctcctgaatttcat |
| *RhaRS-P_RhaBAD_* | TTCGAGCTCGGTACCCGGGGggctctgctgtagtgagtgggttgcgctccggcagcggtcctgatcccccgcagaaaaaaaggatctcaagaagatcctttgatcttttctacggcgcgcccagctgtctagggcggcggatttgtcctactcaggagagcgttcaccgacaaacaacagataaaacgaaaggcccagtctttcgactgagcctttcgttttatttgatgcctttaattaaagcggataacaatttcacacaggaggccgatcgcgtttaatctttctgcgaattgagatgacgccactggctgggcgtcatcccggtttcccgggtaaacaccaccgaaaaatagttactatcttcaaagccacattcggtcgaaatatcactgattaacaggcggctatgctggagaagatattgcgcatgacacactctgacctgccgcagatattgattgatggtcattccagtctgctggcgaaattgctgacgcaaaacgcgctcactgcacgatgcctcatcacaaaatttatccagcgcaaagggacttttcaggctagccgccagccgggtaatcagcttatccagcaacgtttcgctggatgttggcggcaacgaatcactggtgtaacgatggcgattcagcaacatcaccaactgcccgaacagcaactcacccatttcgttagcaaacggcacatgctgactactttcatgctcaagctgaccgataatctgccgcgcctgcgccatcccaacgctacctaagcgccagtgtggttgccctgcgctggcgctaaatcccggaatcgccccctgccagtcaagattcagcttcagccgctctgggcaataaataatattctgcaaaaccagatcgttaacggaagcgtaggagtgtttatcgtcagcatgaatgtaaaagagatcgccacgggtaatgcgataagggcgatcgttgagtacatgcagaccattaccgcgccagacaatcaccagctcacaaaaatcatgtgtatgttcggcaaagacatcttgcggataacggtcagccacagcgactgcctgctggtcgctggcaaaaaaatcatctttgagaagttttaactgatgcaccaccgtggctacctcggccagagaacgaagttgattattcgcaatatggcgtacaaatacgttgagaagattcgcgttattgcagaaaaccatcccgtccctgacgaatatcacgcggtgaccagttaaactctcggcgaaaaagcgtcgaaaagtggttactgtcgctgaatccacagcgataggcgatgtcagtaacgctggcctcactgtggcgtagcagatgtcgggctttcatcagtcgcaggcgattcaggtaacgctgaggcgtcagtcccgtttgctgcttaagctgccgatgtagcgtacgcagtgaaagagaaaattgatccgccacggcatcccaattcacctcatcggcaaaatggtcctccagccaggccagaagcaagttgagacgtgatgcgctgttttccaggttctcctgcaaactgcttttacgcagcaagagcagtaattgcataaacaagatctcgcgactggcggtcgagggtaaatcattttccccttcctgctgttccatctgtgcaaccagctgtcgcacctgctgcaatacgctgtggttaacgcgccagtgagacggatactgcccatccagctcttgtggcagcaactgattcagcccggcgagaaactgaaatcgatccggcgagcgatacagcacattggtcagacacagattatcggtatgttcatacagatgccgatcatgatcgcgtacgaaacagaccgtgccaccggtgatggtatagggctgcccattaaacacatgaatacccgtgccatgttcgacaatcacaatttcatgaaaatcatgatgatgttcaggaaaatccgcctgcgggagccggggttctatcgccacggacacgttaccagacggaaaaaaatccacactatgtaatacggtcatactggcctcctgatgtcgtcaacacggcgaaatagtaatcacgaggtcaggttcttaccttaaattttcgacggaaaaccacgtaaaaaacgtcgatttttcaagatacagcgtgaattttcaggaaatgcggtgagcatcacatcaccacaattcagcaaattgtgaacatcatcacgttcatctttccctggttgccaatggcccattttcctgtcagtaacgagaaggtcgcgaattcaggcgctttttagactggtcgtaatgaaattcAGGAGGTGGTCGACAcatgagcaagggcgaagaac |
| *P_benA_* | ttcgagctcggtacccggggCACCTGGTAGCTGCAAAAGGATTTTTAGAAAAGCGTGAGTTGTTCGAAGCATTGCCATTTTCTGAAGTTACCGAAAAAGTACCGAACATCCGTAAATCTGGATAACGTTCTGCACAATCCGGATAGCCCCCCGCCAGCCGTCTCCCTAACCTGACCAGGTCTAAACAATAACAAGGGAGACCCTGGCCcatgagcaagggcgaagaac |
| *P_pedE_* | ttcgagctcggtacccggggGCAAGCAACACATTGCATTTTGTTGTCATTGTTATGCCCTCACCATTGCACACAAAGCCTCAACCACAGAGGCCGTGCCGCCATCTTAGGAACAGCCCGCGGGCGGCCGAATGCTGCTTTGGTGGCCGGGCCTTAATCCCTTGGTAGTAGGGCTTGTGGCGAAGGGCCAATTGAGGTGCCGCGCGGGAATTATTCCCGGCAAGGGCGGGACTTTTTCCGTATCCCGCGATGGCCGCTGCGCCCCACCCTGTGCAGGCCAAAACCTCCACTGGGAACTGCAACCcatgagcaagggcgaagaac |
| *sfgfp-rrnB* | attcgagctcggtacccgggGGATCCATGAGCAAGGGCGAAGAACTGTTCACCGGTGTGGTACCGATCCTCGTCGAGCTGGACGGTGATGTGAATGGCCACAAGTTTTCGGTGCGTGGCGAGGGTGAAGGTGACGCGACCAACGGCAAGCTGACTTTGAAATTCATCTGCACCACAGGGAAACTGCCGGTGCCGTGGCCGACCCTGGTGACCACCCTGACGTACGGCGTGCAGTGCTTCAGCCGCTATCCTGATCACATGAAGCGGCACGACTTCTTCAAAAGTGCCATGCCCGAGGGGTATGTGCAAGAGCGCACCATCAGCTTCAAGGACGATGGCACCTACAAGACCCGTGCTGAAGTCAAGTTCGAAGGCGACACGCTTGTCAACCGCATCGAGCTCAAGGGGATTGATTTTAAGGAAGACGGCAACATCCTGGGCCATAAACTGGAATACAACTTCAATTCCCATAATGTCTACATCACCGCAGACAAGCAGAAGAACGGCATAAAGGCCAACTTCAAGATCCGCCACAACGTGGAGGACGGTAGCGTGCAACTGGCCGACCATTACCAGCAGAACACTCCAATCGGCGATGGCCCAGTATTGCTGCCCGACAACCACTACCTGTCGACCCAGTCAGTTTTGTCCAAGGACCCGAACGAGAAGCGCGACCACATGGTGCTGCTGGAGTTCGTCACGGCGGCCGGCATTACCCACGGCATGGACGAACTGTACAAATAAtaatctgctgagtgatagactcaaggtcgctcctagcgagtggcctttatgattatcactttacttatgtgatgggaactgccaggcatAAATAAAACGAAAGGCTCAGTCGAAAGACTGGGCCTTTCGTTTTATCTGTTGTTTGTCGGTGAACGCTCTCctctagagtcgacctgcagg |
| *syfp2* | tccactgggaactgcaacccatggtgagcaagggcgaggagctgttcaccggggtggtgcccatcctggtcgagctggacggcgacgtaaacggccacaagttcagcgtgtccggcgagggcgagggcgatgccacctacggcaagctgaccctgaagctgatctgcaccaccggcaagctgcccgtgccctggcccaccctcgtgaccaccctgggctacggcgtgcagtgcttcgcccgctaccccgaccacatgaagcagcacgacttcttcaagtccgccatgcccgaaggctacgtccaggagcgcaccatcttcttcaaggacgacggcaactacaagacccgcgccgaggtgaagttcgagggcgacaccctggtgaaccgcatcgagctgaagggcatcgacttcaaggaggacggcaacatcctggggcacaagctggagtacaactacaacagccacaacgtctatatcaccgccgacaagcagaagaacggcatcaaggccaacttcaagatccgccacaacatcgaggacggcggcgtgcagctcgccgaccactaccagcagaacacccccatcggcgacggccccgtgctgctgcccgacaaccactacctgagctaccagtccaagctgagcaaagaccccaacgagaagcgcgatcacatggtcctgctggagttcgtgaccgccgccgggatcactctcggcatggacgagctgtacaagtaaaaataaaacgaaaggctcag |

**Supplementary Table S3. Vectors used in this work.**

| **Name** | **Relevant features** | **Reference** |
| --- | --- | --- |
| pSEVA231 | SEVA vector;oriT; oriV(pBBR1); MCS; Km^R^ | (Silva-Rocha et al., 2013) |
| pSTDesR·phi15gp16 | Expression of the growth decoupler gp16;  oriT; oriV(RK2); RhaRS/P_RhaBAD_→MCS; Sm^R^/Sp^R^ | (Lammens et al., 2025) |
| pSEVA231·*sfgfp* | pSEVA231 derivative with *sfgfp-rrnB* insertion | This work |
| pSEVA231·*P_RhaBAD_-sfgfp* | pSEVA231·*sfgfp* derivative with *P_RhaBAD_* insertion | This work |
| pSEVA231·*P_benA_-sfgfp* | pSEVA231·*sfgfp* derivative with *P_benA_* insertion | This work |
| pSEVA231·*P_PedE_-syfp2* | pSEVA231·*sfgfp* derivative with *P_pedE_* insertion and *syfp2* exchange with *sfgfp* | This work |

**Supplementary Table S4. Statistical analysis of entropy related to Figure 1.** Showing p-values for the test for normality, equal variance, and equal means. The threshold for the hypothesis was 0.05. A star (*) marks a T-test with equal variance.

| **Shapiro-Wilk test for normality** | |  |  |
| --- | --- | --- | --- |
| **System** | **p-Value** |  |  |
| Sensor – L-Rhamnose | 9.48E-15 |  |  |
| Sensor – L-Rhamnose + Benzoate | 1.44E-7 |  |  |
| Sensor + GP – L-Rhamnose | 3.46E-20 |  |  |
| Sensor + GP – L-Rhamnose + Benzoate | 2.17E-11 |  |  |
| **Levene's test for equal variance** | | | |
| **System** | Sensor  – L-Rhamnose | Sensor – L-Rham-nose + Benzoate | Sensor + GP –  L-Rhamnose |
| Sensor – L-Rhamnose |  |  |  |
| Sensor – L-Rhamnose + Benzoate | 0.00167 |  |  |
| Sensor + GP – L-Rhamnose | 5.21E-09 | 5.01E-09 |  |
| Sensor + GP – L-Rhamnose + Benzoate | 4.02E-14 | 5.04E-12 | 0.12780 |
| **T-test for equal means** | | | |
| **System** | Sensor  – L-Rhamnose | Sensor – L-Rham-nose + Benzoate | Sensor + GP –  L-Rhamnose |
| Sensor – L-Rhamnose |  |  |  |
| Sensor – L-Rhamnose + Benzoate | 2.65E-30 |  |  |
| Sensor + GP – L-Rhamnose | 1.77E-76 | 4.65E-76 |  |
| Sensor + GP – L-Rhamnose + Benzoate | 1.07E-96 | 6.53E-85 | 8.38E-27* |

**Supplementary Table S5. Statistical analysis of entropy related to Figure 2.** Showing p-values for the test for normality, equal variance, and equal means between responsiveness in dependency on whether benzoate was added with L-Rhamnose or not. The threshold for the hypothesis was 0.05.

| **Shapiro-Wilk test for normality** | |
| --- | --- |
| **System** | **p-Value** |
| Without Benzoate | 0.00354 |
| With Benzoate | 0.00122 |
| **Levene's test for equal variance** | |
| **System** | With Benzoate |
| Without Benzoate | 4.53E-08 |
| **T-test for equal means** | |
| **System** | With Benzoate |
| Without Benzoate | 6.04E-21 |

**Supplementary Table S6. Pulse frequencies and times.**

| **Pulse frequency**  **[h^-1^]** | **Pulse Benzoate I**  **[h]** | **Pulse Benzoate II**  **[h]** | **Pulse EG I**  **[h]** | **Pulse EG II**  **[h]** |
| --- | --- | --- | --- | --- |
| $\frac{1}{10}$ | 5.25 | 5.75 | 8.53 | 7.93 |
|  | 15.25 |  | 18.53 | 17.94 |
|  | 25.25 | 25.8 | 28.55 | 27.95 |
| $\frac{1}{5}$ | 35.25 | 35.83 | 30.33 | 37.96 |
|  | 40.25 | 40.85 | 38.13 | 42.97 |
|  | 45.25 | 45.88 | 43.15 | 47.98 |
| $\frac{1}{3}$ | 50.25 | 50.9 | 48.17 | 52.99 |
|  | 53.25 | 53.93 | 53.18 | 56 |
|  | 56.25 | 56.96 | 56.18 | 59.01 |
| $\frac{1}{2}$ | 59.25 | 59.98 | 59.2 | 62.01 |
|  | 61.25 | 62.01 | 62.22 | 64.02 |
|  | 63.25 | 64.03 | 64.23 | 66.03 |
| $\frac{1}{1.5}$ | 65.25 | 66.06 | 66.23 | 68.04 |
|  | 66.75 | 67.58 | 68.25 | 69.55 |
|  | 68.25 | 69.11 | 69.77 | 71.06 |
|  |  |  | 71.29 |  |
| $\frac{1}{1}$ | 69.75 | 70.64 | 72.78 | 72.57 |
|  | 70.75 | 71.66 | 73.8 | 73.58 |
|  | 71.75 | 72.69 | 74.82 | 74.59 |
| $\frac{1}{0.5}$ | 72.75 | 73.71 | 75.83 | 75.6 |
|  | 73.25 | 74.24 | 76.33 | 76.11 |
|  | 73.75 | 74.76 | 76.93 | 76.62 |
|  | 74.25 | 75.29 | 77.45 | 77.13 |
|  | 74.75 | 75.81 | 77.97 | 77.64 |
|  | 75.25 | 76.34 | 78.07 | 78.15 |
|  | 75.75 | 76.87 | 78.18 | 78.66 |
|  |  |  | 78.3 |  |

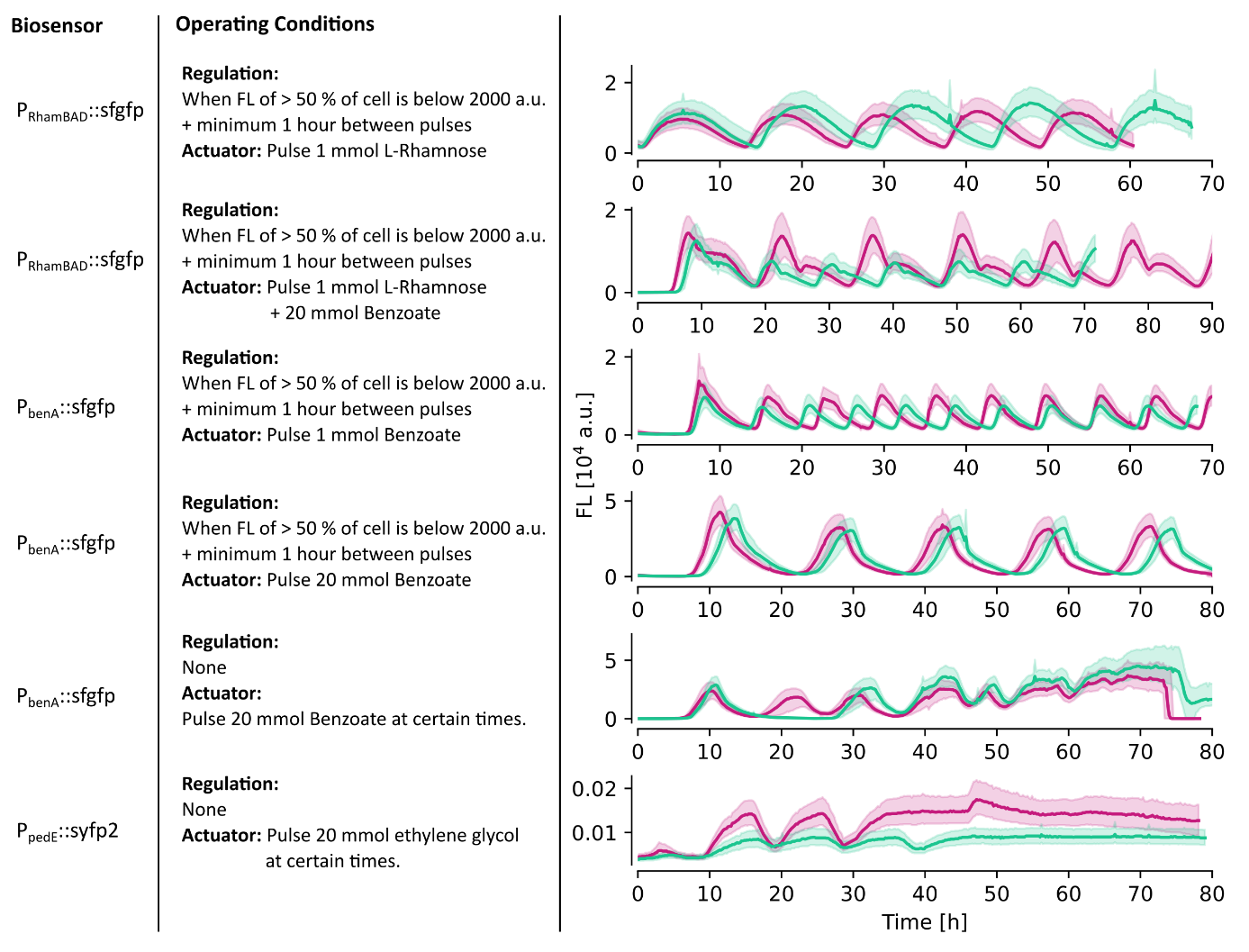

**Supplementary Figure S1**: **Reproducibility of the Segregostat experiments.** Operating conditions and fluorescence evolution of the duplicates in function of biosensors used. The lines represent the median fluorescence, while the shadow shows the interquartile distance.

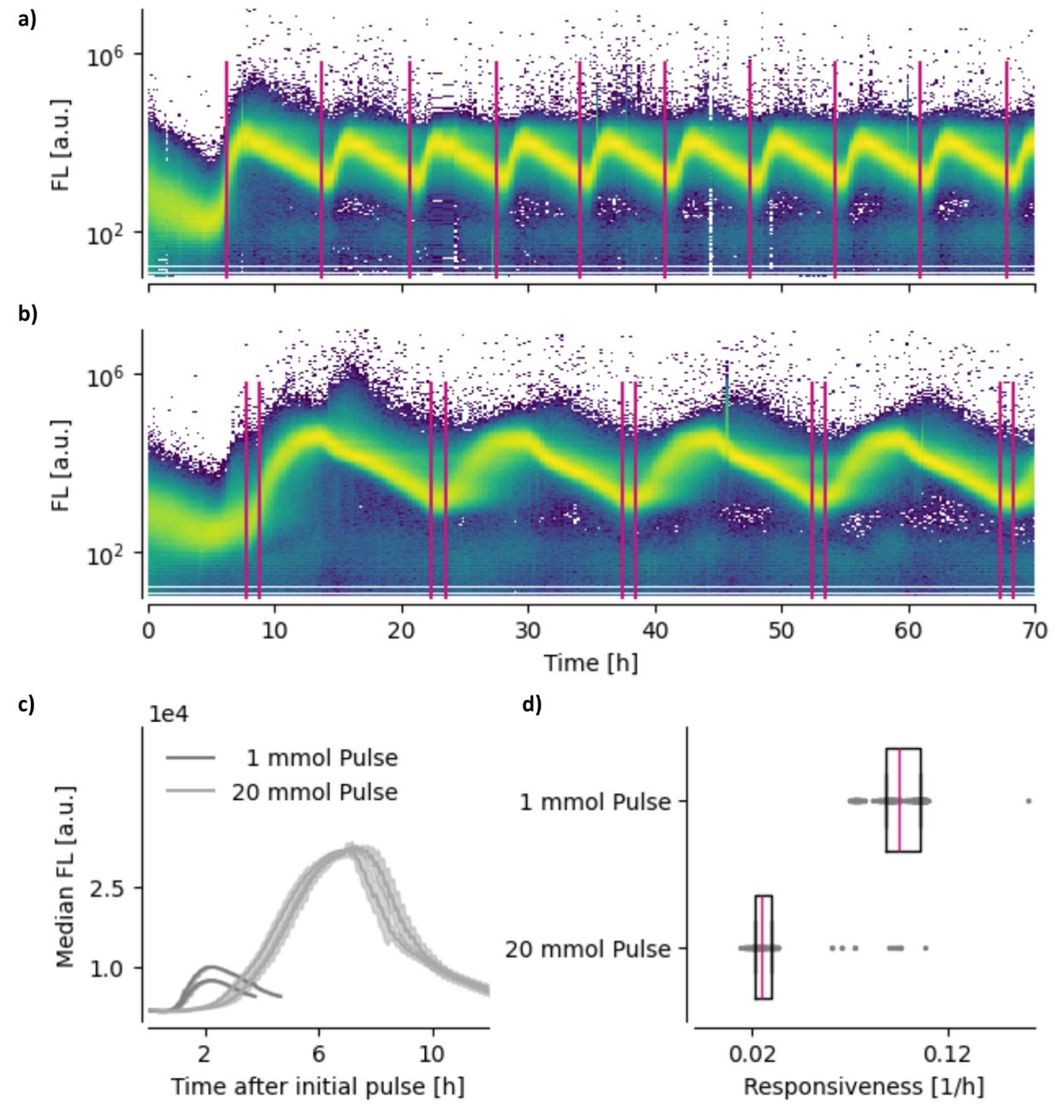

**Supplementary Figure S2:** **Characterization of the activation profile of P. putida KT2440 pSEVA231·P_benA_-sfgfp upon pulsed benzoate feeding**. Fluorescence distributions were monitored using automated flow cytometry during repeated pulses of either 1 mM (a) or 20 mM (b) benzoate per pulse. The regulation activates the benzoate pulse when 50% of the cells emit a fluorescence below 2000 a.u. To prevent overfeeding, a minimum interval of 1 h was maintained between successive pulses. The average median fluorescence (c) and cellular responsiveness (d) illustrate the effect of increased benzoate concentrations on the activation dynamics. Median fluorescence values were averaged after normalizing time to the first pulse of each oscillatory cycle.

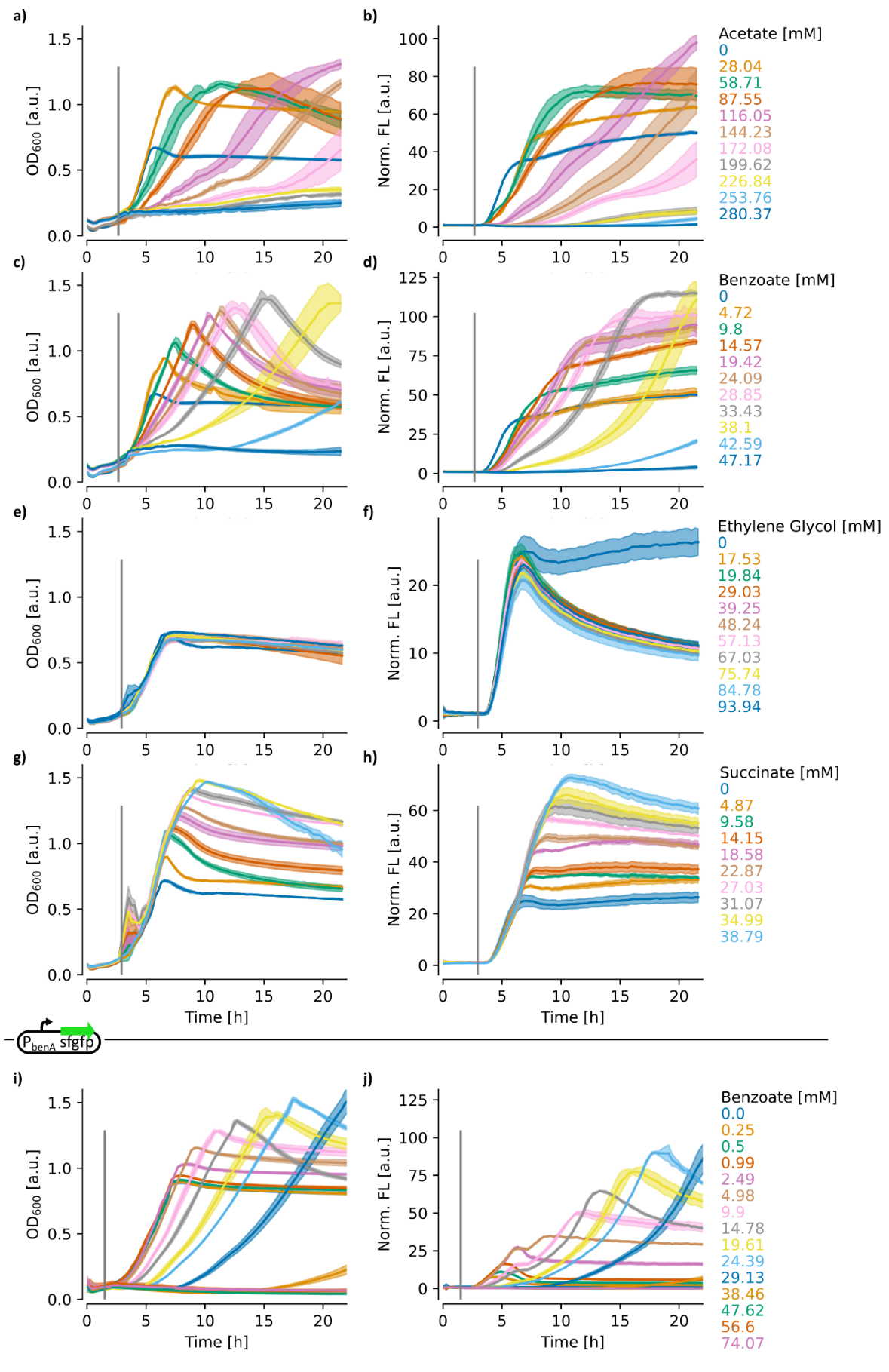

**Supplementary Figure S3:** **Optical density and fluorescence of P. putida pSEVA231·PRhaBAD-sfgfp and P. putida pSEVA231·PbenA-sfgfp under exposure to varying compounds.** Cells were cultivated in 150 µL minimum salt medium supplemented with 2 g/L glucose. When the population reached the exponential growth phase, L-Rhamnose (final concentration 1 mM; 100 mM stock) and various compounds were added (100 or 1000 mM). Development of optical density (OD) and normalised fluorescence is shown for acetate (a, b), benzoate (c, d), ethylene glycol (EG) (e, f), succinate (g, h), and for benzoate with cells containing the PbenA-sfgfp biosensor (I,j). In the latter case, only

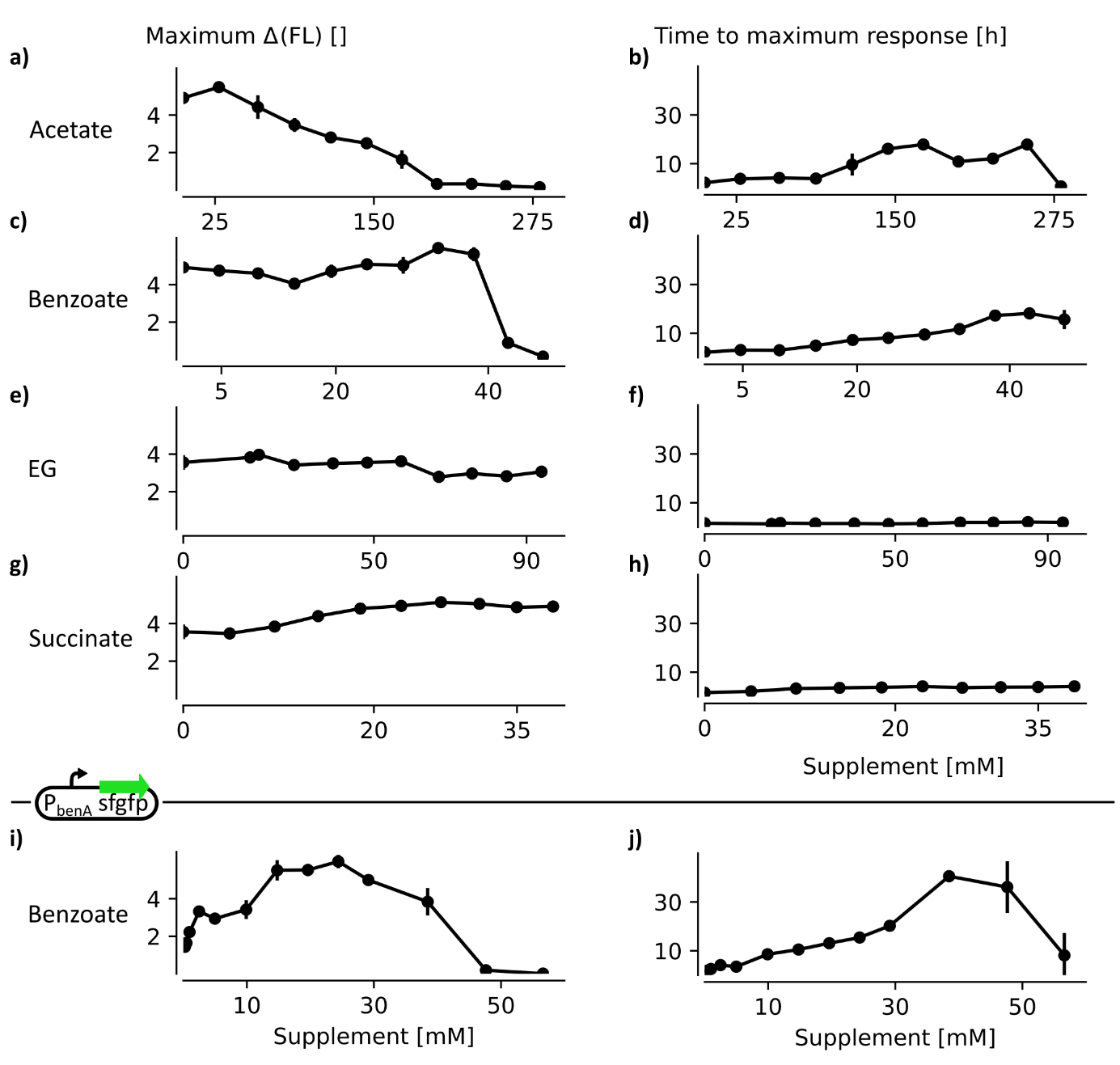

**Supplementary Figure S4: Maximum fluorescence production and the time of maximum production after the pulse in relation to the supplemented compound and compound concentration within the multiplate reader experiments.** P. putida pSEVA231·PRhaBAD-sfgfp in exposure to acetate (a,b), benzoate (c,d), ethylene glycol (EG) (e,f), succinate (g,h) and P. putida pSEVA231·PbenA-sfgfpin exposure to benzoate (I,j). Responsiveness is calculated by dividing maximum fluorescence production by the time of occurrence after the pulse.

**Supplementary Note S1: Resource Allocation Model**

**Model Overview**

A dynamic resource allocation model was developed to describe microbial growth on two carbon sources, glucose (S) and benzoate (B), where benzoate exerts an inhibitory effect on cellular growth and metabolism. The model assumes that total biomass (X) consists of four functional sectors:

$$X=R+C_{S}+C_{B}+T$$

(1)

Where $\left( R \right)$ represents unspecialized biomass that serves as a reservoir for cellular resource allocation, $\left( C_{S} \right)$ represents biomass specialized for glucose utilization, $\left( C_{B} \right)$ for benzoate utilization, respectively, and $\left( T \right)$represents a tolerance sector that mitigates benzoate-induced inhibition. The sectors use [g/L] as the unit for total biomass and represent the amount of biomass allocated to this sector, including protein production and metabolic resources like ATP and reaction equivalents.

The relative abundance of the specialized sectors $\left( C_{S}/X \right),\left( C_{B}/X \right)$and $\left( T/X \right)$ determines substrate uptake capacity and stress tolerance. Cellular resources can be dynamically allocated from the unspecialized sector $\left( R \right)$ toward the specialized sectors in response to environmental conditions.

**Substrate uptake kinetics**

Substrate consumption is described using Monod-type uptake functions modulated by sector allocation and inhibition:

$$\frac{dS}{dt}=v_{s} \frac{\left( C_{s}/X \right)}{K_{S}+\left( C_{s}/X \right)} \frac{S}{K_{S2}+S} i X$$

(2)

$$\frac{dB}{dt}=v_{b} \frac{\left( C_{b}/X \right)}{K_{B}\cdot ii+\left( C_{b}/X \right)} \frac{B}{K_{B2}+B}i X$$

(3)

where $\left( v_{S} \right)$ and $\left( v_{B} \right)$ are the maximum uptake coefficients for glucose and benzoate, respectively.

**Inhibition Functions**

Benzoate inhibition is represented using a uncompetitive inhibition equation. The inhibition effect of benzoate is neutralized by the allocation of biomass to the tolerance sector $\left( T \right)$:

$$i=\frac{1}{1+\frac{B}{K_{i}+\frac{\left( T/X \right)}{K_{iT}}}}$$

(4)

In addition, glucose represses benzoate utilization through the regulatory term:

$$ii=1+\frac{S}{K_{is}}$$

(5)

where $\left( K_{i} \right),\left( K_{i}T \right)$ and $\left( K_{i}s \right)$ are inhibition constants.

**Formation of specialized biomass sectors**

Specialized biomass sectors are generated by allocating resources from the unspecialized biomass pool $\left( R \right)$. Formation rates are described as:

$$r_{Cb}=v_{e}R\left( \frac{B}{K_{cb}+B} \right)i$$

(6)

$$r_{Cs}=v_{e}R\left( \frac{S}{K_{cs}+S} \right)i$$

(7)

$$r_{T}=v_{e}R\left( \frac{(C_{B}/X)}{K_{T}+(C_{B}/X)} \right)i$$

(8)

with $\left( ve \right)$ describing the maximum resource allocation rate and $\left( K_{Cb} \right),\left( K_{Cs} \right)$ and $\left( K_{T} \right)$ are allocation constants. The tolerance sector is induced in response to increasing investment in benzoate-utilizing biomass, reflecting the cellular requirement to mitigate benzoate-associated stress.

The abundance of each specialized sector $P_{i}\left( P_{i}\in C_{s},C_{b},T \right)$ changes according to its formation rate and a degradation process:

$$\frac{dP_{i}}{dt}=r_{i}- deg P_{i}$$

(9)

The degradation coefficient increases with benzoate concentration and is also mitigated by the tolerance sector:

$$deg=d_{0}+d \frac{B}{K_{d}\left( 1+\frac{\left( T/X \right)}{K_{iT}} \right)+B}$$

(10)

here, $\left( d_{0} \right)$ is the basal degradation rate, and d is the maximum benzoate-dependent degradation coefficient.

The unspecialized biomass pool $\left( R \right)$ is replenished through substrate assimilation and depleted by allocation to specialized sectors. Its dynamics are given by:

$$\frac{dR}{dt} =- Y_{R\_S} \frac{dS}{dt} - Y_{R\_B} \frac{dB}{dt} - Y_{R\_Cs} r_{Cs}- Y_{R\_Cb} r_{Cb}- Y_{R\_T} r_{T}$$

(11)

Where the yield coefficients $\left( Y \right)$ define the amount of unspecialized biomass generated from substrate consumption or required for the formation of specialized biomass sectors.

**Parameter Estimation and Model Validation**

Model parameters were estimated using the experimental dataset presented in **Supplementary Fig S3i**, comprising 45 growth curves obtained under 15 different benzoate pulse concentrations. Optical density (OD) measurements were converted to biomass concentrations using the conversion factor of 0.7. Data until the end of the exponential growth phase were included in the fitting procedure.

To minimize overfitting, the dataset was randomly partitioned into training and validation sets. Parameter estimation was performed on 80% of the growth curves, with the remaining 20% reserved for independent model validation.

Optimization was conducted using the Differential Evolution algorithm implemented in the SciPy Python library. Differential Evolution was selected due to its robustness in identifying global optima in high-dimensional, nonlinear parameter spaces. The optimization procedure consisted of two stages, a global and sequential fine fitting.

For each optimization, a population size of 60 candidate solutions was evolved over 1000 generations. The objective function minimizes the discrepancy between the model and the datasets by calculating the mean of the mean squared percentage error (MSPE) across all growth curves *m*.

$$MMSPE = \frac{1}{m} \sum\frac{1}{n}\sum\frac{\left( y_{data}-y_{model} \right)^{2}}{{y_{data}}^{2}}$$

(12)

**Supplementary Table S6. List of parameters**

| Variable | Unit | Initial Value |  |  |
| --- | --- | --- | --- | --- |
| B  S  R  Cb  Cs  T  X | g/L  g/L  g/L  g/L  g/L  g/L  g/L | 0  2  0  0  0.056  0  0.056 |  |  |
| Parameter | Unit | Boundaries | 1^st^ Fit | 2^nd^ Fit |
| vb  vs  ve  K_B_  K_B2_  K_S_  K_S2_  K_Cb_  K_Cs_  K_T_  Y_R_S_  Y_R_B_  Y_R_Cs_  Y_R_Cb_  Y_R_T_  K_i_  K_iT_  K_iS_  d_0_  d  K_d_ | 1/h  1/h  1/h  gL/gL  g/L  gL/gL  g/L  g/L  g/L  gL/gL  gL/gL  gL/gL  gL/gL  gL/gL  gL/gL  g/L  1/ gL^-1^  g/L  1/h  1/(hgL^-1^)  - | [1,5]  [1,5]  [0.01,10]  [0.001,0.1]  [0.00001,0.1]  [0.001,0.1]  [0.00001,0.1]  [0.0001,1]  [0.0001,1]  [0.001,10]  [0.3,0.8]  [0.3,0.8]  [1.1,10]  [1.1,10]  [1.1,10]  [0.0001,1]  [0.0001,1]  [0.0001,100]  [0.00001,1]  [0.01,10]  [0.1,100] | 4.376  1.0173  8.4769  0.0511  0.0274  0.0998  0.021  0.3771  0.0549  7.8695  0.7539  0.453  5.9497  2.2826  2.5549  0.9883  0.9701  53.3898  0.0061  1.7774  86.759 | 4.994  1.0  9.1367  0.0013  0.00001  0.1  0.0775  0.0889  0.1925  6.7421  0.3795  0.4512  9.6996  1.1027  1.1924  0.993  0.8454  33.9116  0.0  1.6933  74.9608 |

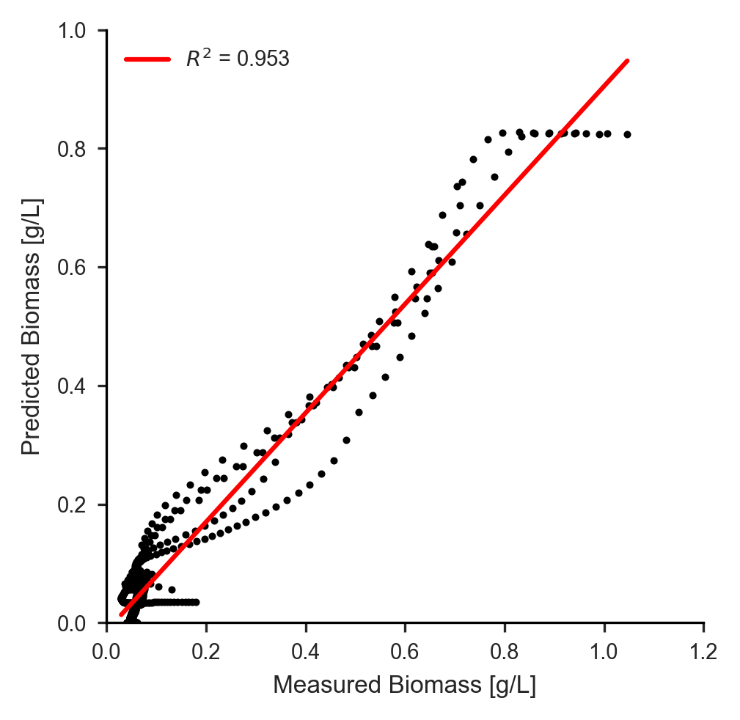

**Supplementary Figure S4:** Verification of the resource allocation model by correlating measured and predicted biomass of all verification data sets. The verification was performed with 20% of the datasets including 8 growth curves.

.

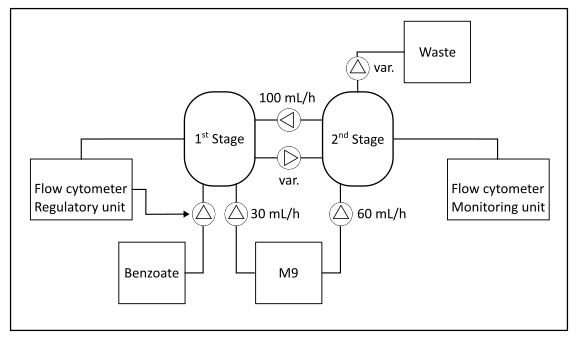

**Supplementary Figure S5:** Schematic of the connected bioreactor system. Both bioreactors receive minimal-salt media at different rates. The media contains 1 g/L glucose and 1 mM benzoate as carbon sources. Additional benzoate is pulsed under FC control. If no collapse is detected, 5 mmol of benzoate (1 M stock) is pulsed after each measurement (~20 mL/h). Variable efflux rates are enabled by an overflow standpipe connected to a pump set to 250 mL/h, thereby maintaining reactor volume.

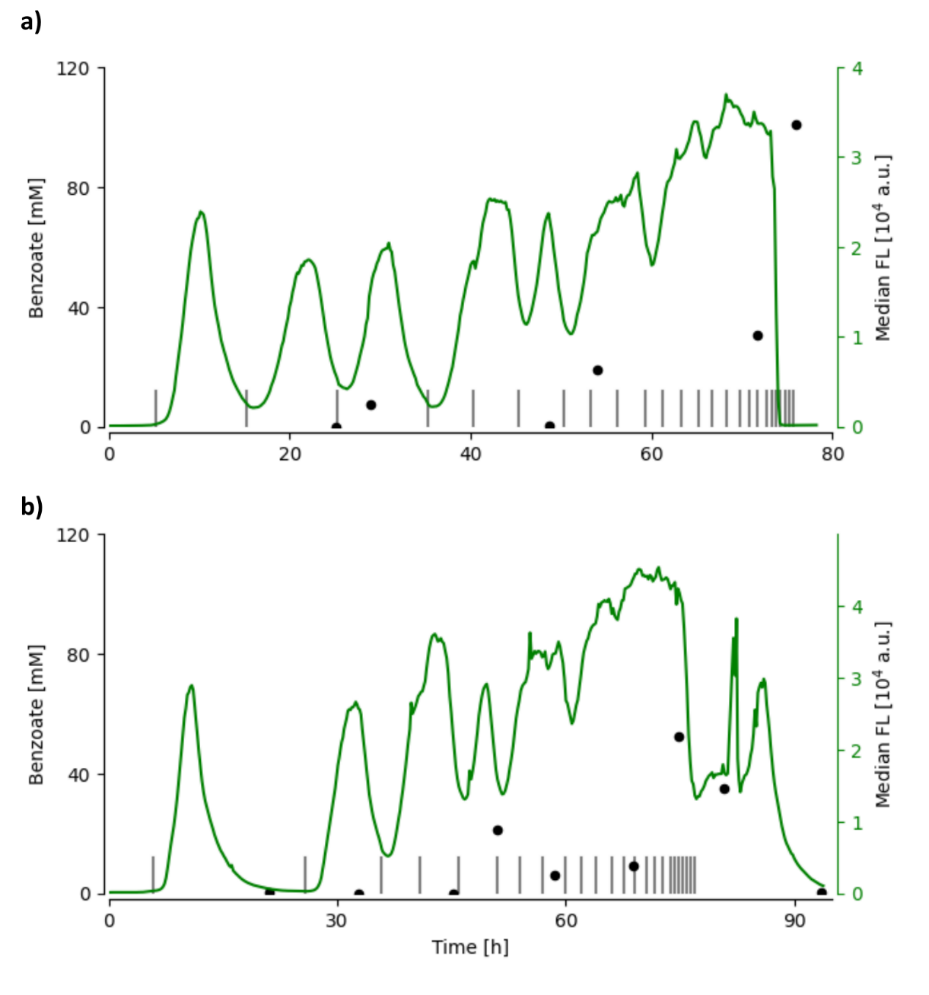

**Supplementary Figure S6:** **Median fluorescence and benzoate concentration while conducting increased pulse frequencies of benzoate upon P. putida pSEVA231·P_benA_-sfgfp.** The addition of 20 mmol benzoate is represented by grey vertical lines. The two panels show the data acquired over technical replicates.

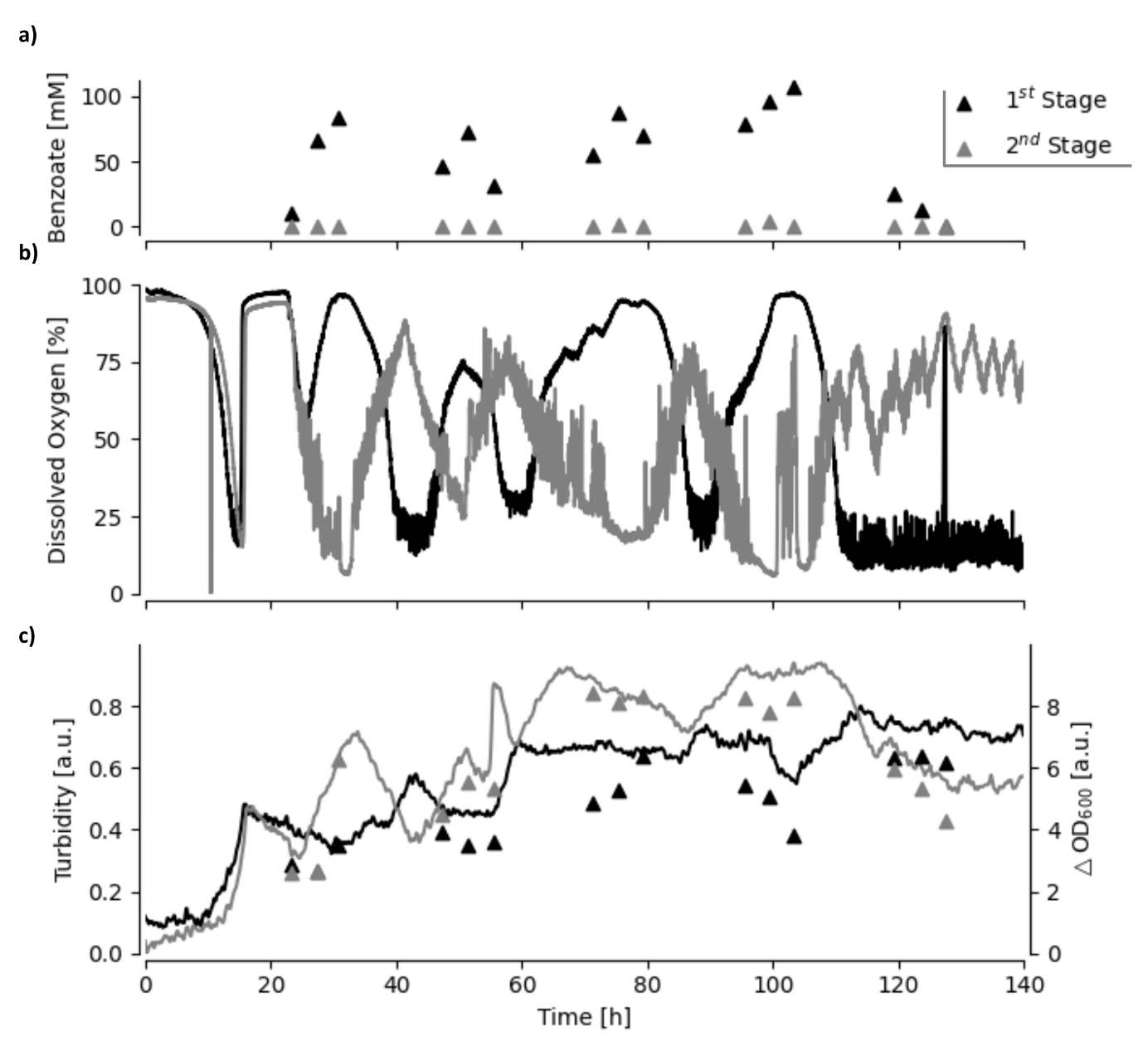

**Supplementary Figure S7:** **Benzoate (a), dissolved oxygen (b), online turbidity and offline OD600 (c) within the coupled bioreactor.** Benzoate addition was controlled by the autonomous flow cytometer on the first stage. As long as less then 10 % of cells exhibit a fluorescence below 2000 a.u., 5 mmol benzoate were pulsed.
